# Prevalence of and gene regulatory constraints on transcriptional adaptation in single cells

**DOI:** 10.1101/2023.08.14.553318

**Authors:** Ian A. Mellis, Nicholas Bodkin, Madeline E. Melzer, Yogesh Goyal

**Affiliations:** Department of Pathology and Cell Biology, Columbia University Vagelos College of Physicians and Surgeons, New York, NY, USA; Aaron Diamond AIDS Research Center, Columbia University Vagelos College of Physicians and Surgeons, New York, NY, USA; Department of Cell and Developmental Biology, Feinberg School of Medicine, Northwestern University, Chicago, IL, USA; Center for Synthetic Biology, Northwestern University, Chicago, IL, USA; Robert H. Lurie Comprehensive Cancer Center, Feinberg School of Medicine, Northwestern University, Chicago, IL, USA

## Abstract

Cells and tissues have a remarkable ability to adapt to genetic perturbations via a variety of molecular mechanisms. Nonsense-induced transcriptional compensation, a form of transcriptional adaptation, has recently emerged as one such mechanism, in which nonsense mutations in a gene can trigger upregulation of related genes, possibly conferring robustness at cellular and organismal levels. However, beyond a handful of developmental contexts and curated sets of genes, to date, no comprehensive genome-wide investigation of this behavior has been undertaken for mammalian cell types and contexts. Moreover, how the regulatory-level effects of inherently stochastic compensatory gene networks contribute to phenotypic penetrance in single cells remains unclear. Here we combine computational analysis of existing datasets with stochastic mathematical modeling and machine learning to uncover the widespread prevalence of transcriptional adaptation in mammalian systems and the diverse single-cell manifestations of minimal compensatory gene networks. Regulon gene expression analysis of a pooled single-cell genetic perturbation dataset recapitulates important model predictions. Our integrative approach uncovers several putative hits—genes demonstrating possible transcriptional adaptation—to follow up on experimentally, and provides a formal quantitative framework to test and refine models of transcriptional adaptation.

## Introduction

Cells have a remarkable ability to sense changing conditions and exhibit robustness in response to perturbations ^1–9^. Several mechanisms underpinning this plasticity have been proposed and more continue to be reported. These mechanisms—operational at multiple levels of biological organization—include protein feedback loops, adaptive mutations, and single-cell molecular variabilities ^10–16^. One recently reported robustness mechanism is a type of transcriptional adaptation, wherein nonsense mutations, i.e., mutations that result in premature termination of protein synthesis, can remarkably trigger the transcription of related genes, including paralogs ^14,15,17–19^. This adaptation, known as nonsense-induced transcriptional compensation, can enable cells and tissues to escape otherwise fatal mutations and function normally ^14,15^. Besides revealing a new kind of transcriptional regulation, this discovery proposed a resolution of the long-standing discrepancy in phenotypes between many knockdowns and knockouts of the same gene ^2^.

Mechanisms underlying nonsense-induced transcriptional compensation have been studied for a handful of curated genes in select developmental contexts, particularly in vertebrates ^18–24^. For example, premature termination codons in gene *egfl7* in zebrafish caused upregulation of the Emilin gene family via protein Upf1, resulting in a near-absence of vascular defects in zebrafish ^22^. However, it is unclear if this compensatory behavior is pervasive in other genes, species, and contexts. While some studies have indicated the absence of this kind of compensation in organisms such as yeast ^25,26^, to date, no genome-wide investigation of this behavior has been undertaken for different mammalian cell types and contexts. As a result, several questions remain unanswered. For example, is this adaptive behavior limited to certain genes associated with specific signaling pathways? Similarly, do such compensating gene families tend to be functionally similar, e.g., transcription factors, cytoskeleton molecules, or enzymes? Moreover, is this behavior intrinsic to a gene or dependent on its extrinsic environment, i.e., cell type or local regulation? Furthermore, how prevalent is this behavior across mammalian systems and contexts, e.g., cancer or differentiation? One hypothesis is that nonsense-induced transcriptional compensation occurs only in very specific biological circumstances and organisms. Alternatively, it is possible that the transcriptional adaptation as a mode of cellular robustness is ubiquitous and occurs more commonly than hitherto appreciated. Both of these scenarios have different implications. For instance, the latter hypothesis implies that functional genetic screens for phenotypic outcomes will need to account for transcriptional adaptation. Computational analysis of existing datasets can lead the way in resolving these alternatives, which can subsequently be tested experimentally.

Another set of questions center around the regulatory constraints on upregulated paralogs and their downstream effector molecules ^27,28^. In particular, nonsense-induced transcriptional compensation can result in incomplete phenotypic penetrance, such that there are either attenuated defects or a subset of cells or organisms which continue to have strong defects despite compensation. In some cases, compensation can happen without necessarily rescuing a phenotypic defect induced by knockout mutations ^24,29–31^. These observations, coupled with the documented evidence that transcription is bursty ^32^, raise the possibility that inherent stochasticity underlying the compensatory gene regulatory networks may translate into single-cell differences. Single-cell differences, in turn, could result in incomplete penetrance, particularly for phenotypes resulting from variable downstream effects on relevant effector gene expression. However, precisely how compensated expression fluctuations manifest into downstream effects has not been formally investigated. For example, what is the ensemble of gene expression distributions of effector molecules post-compensation? Under what conditions can we expect the system to exhibit robustness of the distribution shape and mean? In a similar vein, does the answer depend on the nature of interactions or network size (negative or positive; one or multiple paralogs)? Resolving these single-cell possibilities with theoretical formulations can provide a mechanistic basis for the observed phenotypic penetrance, and aid in the design of predictive experiments, especially as single-cell technologies become more accessible ^33,34^.

Here we combine computational analysis of existing datasets with mathematical modeling of stochastic gene regulatory interactions to address the questions posed above. First, we argue that a systematic bioinformatic analysis of publicly available transcriptome-wide datasets that rely on CRISPR-Cas9-mediated mutagenesis can, in principle, suggest the presence of transcriptional compensation, or lack thereof. Indeed, our unbiased computational pipeline surveying over 200 publicly available datasets not only recovers the known and validated gene targets that display transcriptional compensation, but also reveals the breadth of transcriptional compensation across mammalian cell types and biological processes. Second, we develop stochastic mathematical models of biallelic gene regulation and simulate over tens of millions of cells. We find that even a relatively parsimonious model of transcriptional adaptation can lead to diverse population-level gene expression distributions of downstream effectors, some of which we recapitulate in our regulon robustness analysis of experimental single-cell sequencing datasets. Our integrative framework is generalizable and lays the foundation for future work to test our findings experimentally and to refine models of transcriptional compensation.

## Results

### A generalizable framework for analyzing CRISPR-Cas9 knockouts paired with RNA-sequencing reveals upregulation of knockout-target paralogs

We wondered whether transcriptional adaptation to mutation—specifically nonsense-induced transcriptional compensation—is widespread in vertebrates, both across genes and biological contexts. To address this question, we took advantage of a feature common in published experimental designs: CRISPR-Cas9-based knockout engineering. When paired with a guide RNA, Cas9 creates a double-stranded DNA break at a predefined site in a target gene, after which endogenous non-homologous end joining repair processes induce a random insertion-deletion (indel) mutation ^35–38^. On average, two-thirds of indels in coding regions will induce a frameshift by random chance. In turn, this frameshift will render the resulting open reading frame of the mutant different from the wild-type and cause a premature termination codon ^39^. Since nonsense-induced transcriptional compensation is proposed to occur as a result of premature termination codons, we hypothesized that transcriptomic data from Cas9-based knockout experiments could reveal the presence—or absence—of potential nonsense-induced transcriptional compensation (Figure 1A). Further, even if a specific nonsense or frameshift allele is not isolated and expanded (i.e., the mutated population is polyclonal), at least two-thirds of Cas9-affected alleles in Cas9-treated cells will be nonsense mutants.

**Figure 1:**
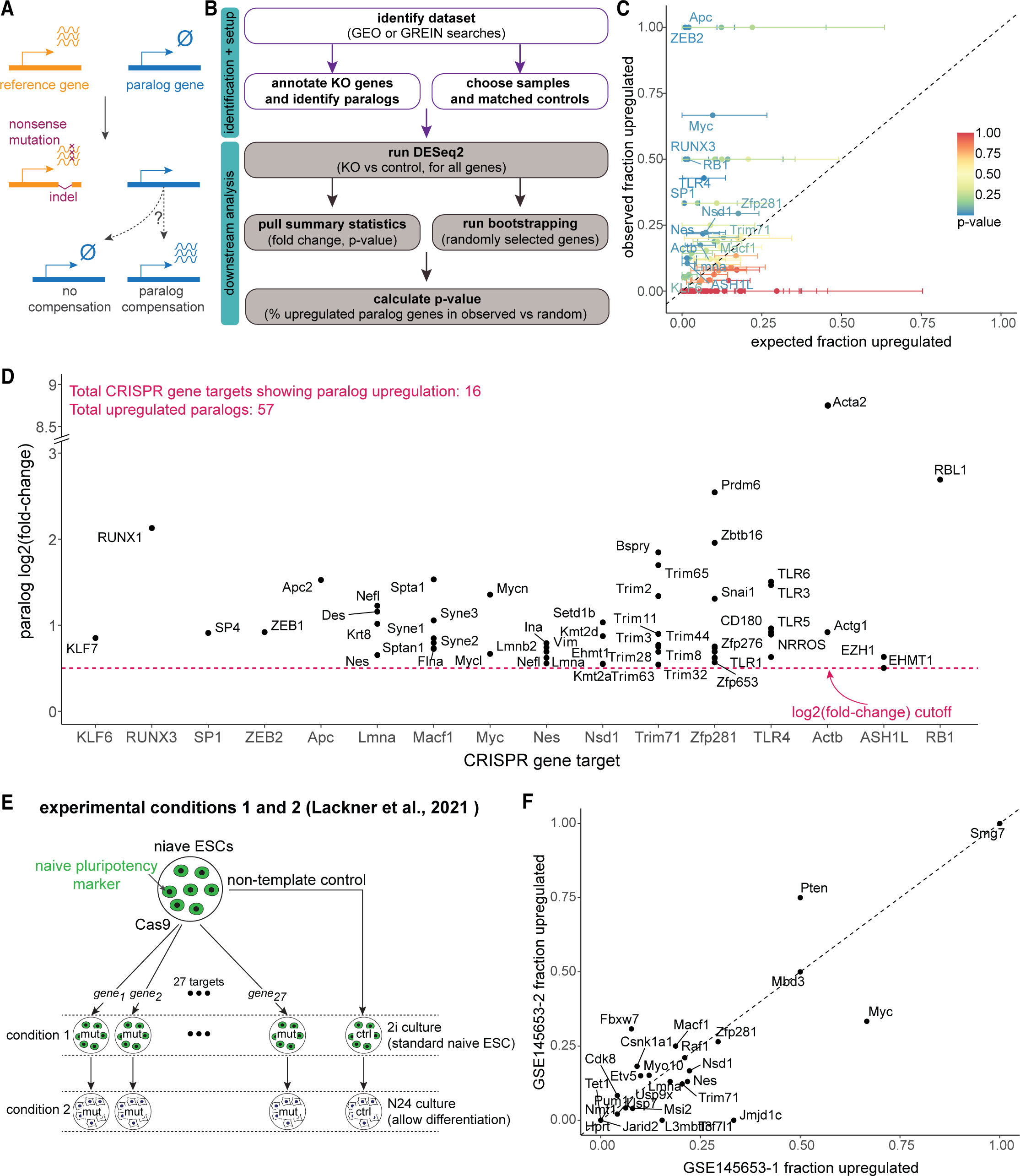
Inferring prevalence of transcriptional adaptation in transcriptomic datasets A. Schematic of transcriptional adaptation after Cas9-mediated mutagenesis. After indel mutation, frameshifts often occur, which lead to premature termination codon formation and resultant nonsense-mediated decay. We hypothesized that paralogs may be upregulated in genes with nonsense-induced transcriptional compensation-type transcriptional adaptation. B. Schematic of analytical workflow. We mined publicly available RNA-seq datasets for differential expression of paralogs of CRISPR/Cas9 knockout targets after mutation. Randomly selected bootstrap sampled genes are chosen to have similar average expression levels as respective paralogs of interest. C. Per-knockout-target paralog differential expression results. Per knockout target: expected frequency of paralog upregulation based on bootstrap analysis on the x axis, observed frequency of paralog upregulation on the y axis. Differential expression counted if log2 fold-change > 0.5 and adjusted p-value < 0.05. Error bars represent the standard error of the null expected mean frequency of paralog upregulation across 10000 bootstrap samples (see Methods). D. Per-upregulated-paralog differential expression magnitude. For significantly upregulated paralogs of knockout targets showing transcriptional adaptation in (C), log_2_ fold-change relative to controls. Knockout targets on x axis in arbitrary order. E. Schematic of experimental design for analysis of paralog upregulation frequency for 27 knockout targets in mouse naive embryonic stem cells (ESC), adapted from Lackner et al., 2021. F. Comparison of paralog upregulation frequency in 2i (standard naive ESC culture condition) versus N24 (removal of 2i for 24 hours, allowing exit from naive pluripotency) for 27 targets.

Since nonsense-induced transcriptional compensation can depend on sequence homology ^14,15^, we first developed a robust methodology for choosing genes that may compensate for a knockout target. There are several documented methods for choosing related, potentially compensating, genes. These range from considering whole protein families to identifying more recently ancestrally related paralog genes to performing genome-wide local alignment searches ^14,22^. We decided to use paralog genes in our analysis as they are consistently annotated and identifying Ensembl-annotated paralogs does not depend on individually optimized local alignment search parameters ^40^. We then performed a comprehensive literature search for published, publicly available datasets for CRISPR-Cas9 knockout experiments paired with bulk RNA sequencing of both nontemplate controls and knockout target samples. We analyzed mouse and human samples. Furthermore, we prioritized published datasets that included multiple parallel knockout experiments. In total, we screened over 200 datasets in the NIH’s Gene Expression Omnibus (GEO), and identified 36 GEO entries with a total of 220 initially analyzable CRISPR gene targets meeting our experimental design criteria, including 76 in mice, 144 in humans (Figure 1B, Table S1). After quality control, paralog lookup, and differential expression filters, we proceeded to analyze a total of 74 gene targets and their respective nontemplate controls (see Methods). The datasets analyzed include knockouts engineered for the study of a variety of biological phenomena, including organ development, reprogramming to pluripotency, and tumor responses to targeted therapies, among others (Table S1).

With our collection of quality-controlled datasets, we examined whether nonsense-induced transcriptional compensation may exist more widely than previously reported. Specifically, we asked whether paralogs of knockout targets were upregulated after knockout more frequently than would be expected by random chance (see Methods). We found that 16 out of 74 knockout targets had significant upregulation of their paralogs (Figure 1C,D, Table S2). We confirmed this result was not specific to our thresholds for differential expression (adj. P-value < 0.05 and log_2_(fold change) > 0.5) by repeating the analysis using other differential expression or average expression paralog inclusion criteria (Figure S1A,B). Gene hits include *ASHL1, KLF6, RB1, RUNX3, SP1, TLR4,* and *ZEB2* in humans and *Actb, Apc, Lmna, Macf1, Myc, Nes, Nsd1, Trim71,* and *Zfp281* in mice. Our analysis is largely consistent with the published findings of related-gene upregulation after nonsense mutation of 3 target genes found to demonstrate nonsense-induced transcriptional compensation by El-Brolosy et al., 2019 (*Fermt2*, *Actg1*, *Actb*; Figure S1C). It is also important to note that our work recapitulated these earlier findings despite using different related-gene-inclusion criteria (local alignment in *El-Brolosy et al., 2019*, vs. paralog identity here). Further, while we observed some degree of transcriptional upregulation in paralogs of all 3 target genes, only *Actb* was deemed significant by our bootstrap analysis pipeline. This result suggests that beyond the 16 significant hits reported in our study, our paralog-based analysis is relatively more stringent, perhaps leading to false-negative findings. Remarkably, there were 2 CRISPR targets in our dataset that are paralogs of each other, *Lmna* and *Nes*, and we found that both were classified as hits. Moreover, for both *Lmna* and *Nes*, their mutual paralog gene *Nefl* was upregulated upon mutation of either *Lmna* or *Nes* (Figure 1D).

Therefore, despite conservative cutoffs, the shared upregulation of compensating, mutually paralogous genes across independent experiments illustrates the power of our approach and reliability of our findings.

We wanted to check whether paralog upregulation frequencies were consistent across conditions for the same genes. Therefore, we compared paralog upregulation frequency for all 27 knockout targets that met our inclusion criteria for analysis across two conditions in a dataset published in *Lackner et al., 2021*^41^ (see Methods). Lackner and colleagues performed RNA-sequencing on different knockout mouse embryonic stem cell lines, both under standard naive stem cell culture conditions and after a day of differentiation (Figure 1E). In our main analysis (Figure 1C,D), we considered the results from *Lackner et al., 2021* under standard culture conditions to reduce potential confounding effects on expression changes associated with possible divergent differentiation outcomes in each knockout compared against differentiated non-template control lines. Nonetheless, we asked whether there is any agreement in paralog upregulation frequency across the standard undifferentiated and differentiated conditions. We found that for each knockout target, the fraction of paralogs upregulated in standard culture conditions was broadly correlated with the fraction of paralogs upregulated after a day of differentiation (Figure 1E,F; r = 0.832). Therefore, for the tested targets in mouse embryonic stem cells, paralog upregulation frequency is similar in two different conditions, further demonstrating the robustness of our approach.

### Paralog upregulation is also observed in large-scale pooled single-cell CRISPR screens

Next, we wondered whether we could identify additional genes that may exhibit nonsense-induced transcriptional compensation in large pooled knockout experiments with single-cell resolution.

Perturb-seq, CROP-seq, CRISP-seq, and related methods enable pooled parallel single-cell gene expression profiling of dozens or hundreds of knockout targets ^42–44^. Here again, we reasoned that single-cell pooled datasets utilizing Cas9 or equivalent perturbation tools would cause indel mutations in the coding regions of the genes of interest. In addition to the benefits of single-cell resolution of gene expression and its high-throughput, Perturb-seq-style data offers consistency by using a common internal set of non-template-control-treated cells as a comparison for all knockout targets. We identified a large, quality-controlled, pre-processed Perturb-seq dataset with ∼750 distinct guide RNAs using Cas9 as the knockout effector in patient-derived cancer cells. The dataset includes dozens of non-template-control guides and gene-targeting guides directed at >200 target human genes with a wide variety of molecular functions, chosen due to their involvement with cell-intrinsic therapy resistance ^45^. After quality controls, we considered cells representing 143 target genes with 429 total targeting guides as well as 37 non-template-control guides in the main analysis (Figure 2A).

**Figure 2:**
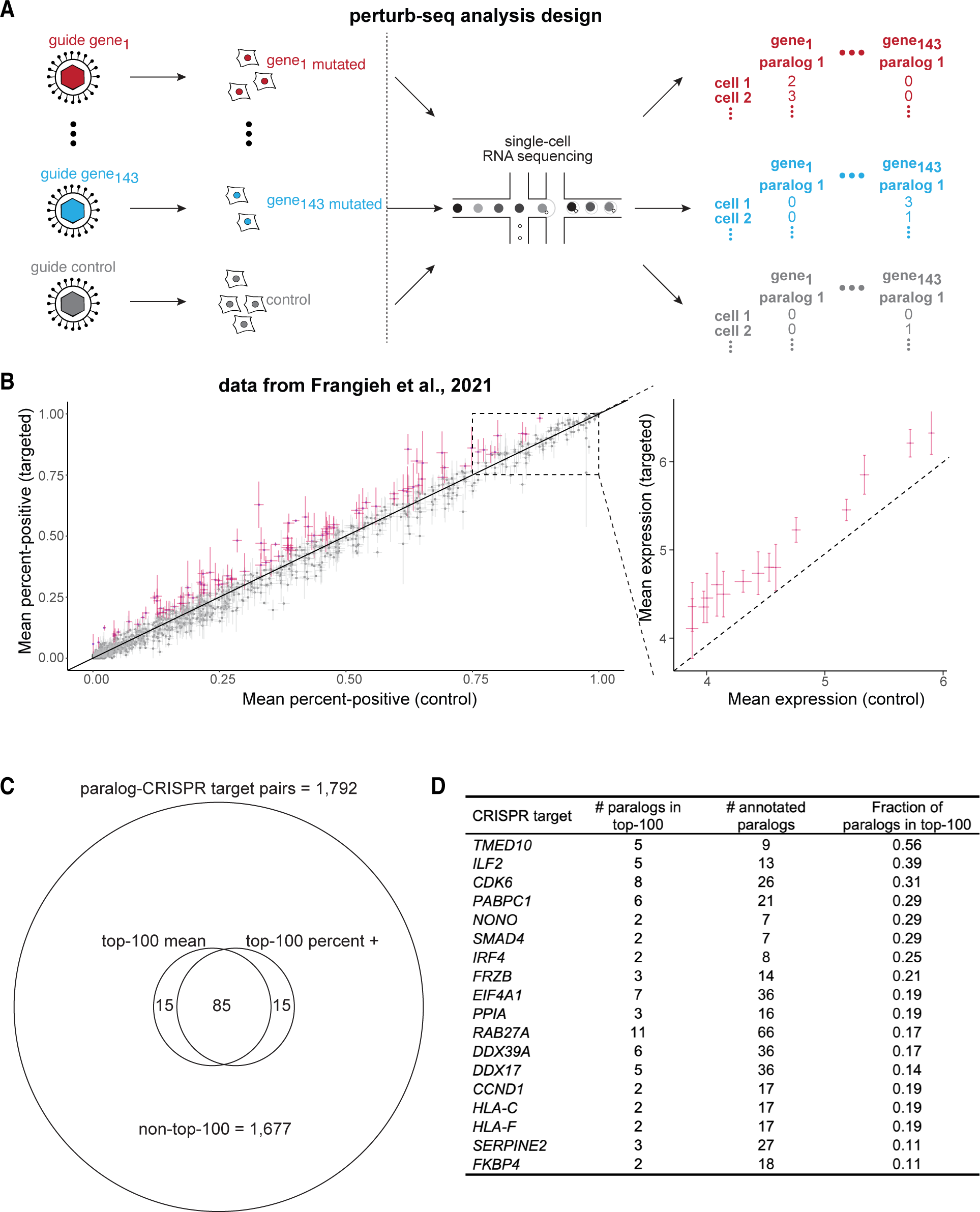
Inferring prevalence of transcriptional adaptation at single-cell resolution in Perturb-seq data A. Schematic of experimental design and paralog gene expression analysis, adapted from Frangieh et al., 2021. 143 gene targets, with a consistent batch of non-template control-treated cells, passing quality control filters (see Methods). B. Perturb-seq-based single-cell paralog expression change after reference gene knockout. Per paralog, percentage of cells positive for that paralog’s expression in non-template control guide-treated cells on x-axis, percentage of cells positive for that paralog’s expression in cells treated with guides targeting a reference CRISPR target for which the gene is a paralog. For paralog genes with percent-positive > 75% in non-template controls, mean expression plotted in inset at right. Paralogs ranked in the top-100 of absolute increases per quantification method marked in magenta. All paralogs of all knockouts shown meeting minimum cell count and UMI count in gray, inclusion criteria listed in Methods. C. Venn diagram summarizing the number of paralog-CRISPR target pairs with the top-100 largest increases in mean expression and/or percent-positive expression levels. 1,792 total paralog-target pairs, of which 1,677 are not in the top-100 largest increase lists. 85 paralog-target pairs in the top-100 largest difference list by both mean and percent-positive analysis. D. List of CRISPR targets with multiple paralogs demonstrating large increases in paralog expression. All CRISPR targets with more than 1 paralog in the top-100 list (see Methods) and with at least 10% of all annotated paralogs in the top-100 lists. A complete list of CRISPR targets with paralogs in the top-100 list is in Table S3.

We then asked whether there was a trend toward increased expression at the single-cell level of any paralogs of each knockout target when compared against non-template-controls. Due to known drop-out events in single-cell RNA-sequencing, we initially focused our analysis on simply counting the fraction of cells with non-zero expression (i.e., “percent positive”) of each paralog of a knockout target. We compared the percent positive cells treated with a targeting guide against cells treated with a control guide ^46,47^. For paralogs with a high baseline expression of at least 75% in control cells, we compared average expression levels instead of percent positive values. In general, the percent positives (and the means) in targeted cells and in controls were well correlated (Figure 2B,C).

Nonetheless, there were many paralog genes in which expression levels may have differed. We rank-ordered the differences between targeted and control cells for all 1,792 paralog-target gene pairs and highlighted the paralogs with the 100 largest absolute increases in percent positive values after respective target knockout (or highest mean increases for those highly expressed at baseline). There were 43 distinct knockout targets with top-100-increase paralogs, including transcription factors,

RNA-binding proteins, signaling pathway components, cell cycle regulators, and cell surface proteins (Figure 2D, Table S3). Additionally, we did a secondary analysis on the subset of gene targets that did not pass our conservative total cell filtering yet showed potential paralog upregulation (Figure S1D).

Particularly, 6 of the 23 paralogs of *TUBB* (Tubulin beta class I) may have been upregulated after *TUBB* mutation (Figure S1D). Intriguingly, others have experimentally shown that mutations of tubulins can lead to tubulin paralog upregulation in a mouse model of neurodegeneration^29^.

We next questioned whether any specific annotated biological pathways, molecular functions, or cellular components were enriched for genes that display transcriptional adaptation by paralogs, either in bulk or in single-cell datasets. Therefore, we conducted gene set enrichment analysis, comparing human knockout targets with significantly upregulated paralogs in bulk data or any paralog in the top-100-increase sets of single-cell data against the full set of human knockout targets tested (see Methods) ^48^. No gene sets were overrepresented among the knockout targets with paralog upregulation. Therefore, transcriptional adaptation, at least for the targets tested, may be broadly acting across the genome and not strongly correlated with functionally defined gene sets or particular signaling pathways.

We also wondered whether higher expression of any genes implicated in the proposed mechanisms of transcriptional adaptation were associated with targets exhibiting paralog upregulation. Therefore, from the two largest datasets, one in human and one in mouse, we extracted the expression levels of 12 genes associated with transcriptional adaptation (members of the COMPASS complex and Upf genes important for nonsense-mediated decay)^14,15,41,49–52^. We compared these 12 genes’ expression levels for each knockout-control pairing in the dataset, grouped by whether the knockout target displayed paralog upregulation. We did not observe any major differences in the 12 genes’ expression levels between targets with signs of transcriptional adaptation versus those without, in either humans or mice (Figure S2). Note, however, that the analysis was limited by low numbers of knockout targets, particularly in the groups displaying paralog upregulation. Future high-throughput studies with more robust datasets will play a critical role in clarifying the role of COMPASS complex component or Upf gene expression levels in transcriptional adaptation.

### Knockout target and paralog features are not associated with paralog upregulation

Since we observed variability in upregulation of different paralogs for a target gene upon CRISPR mutation, we wondered whether paralog-intrinsic factors might be associated with whether a given paralog participates in transcriptional adaptation. Specifically, recent studies have indicated that genes that have some degree of local sequence homology with a nonsense-mutated gene are more likely to be upregulated after mutation, but this has not been systematically investigated ^14^. We checked whether paralog upregulation has any association with the degree of sequence homology (i.e., transcript-wide percent homology) with the knockout target. We found no significant correlation between the degree of sequence homology and the expression change after mutation for bulk CRISPR targets demonstrating transcriptional adaptation (Figure S3A). Next, given the requirement for degraded transcripts from mutated genes in proposed mechanisms of transcriptional adaptation, we wondered whether longer knockout target genes might more often have paralog upregulation. We found no significant correlation between length of gene and the paralog upregulation frequency (Figure S3B). Taken together, our results suggest that nonsense-induced transcriptional compensation may exist for several vertebrate genes, and the paralog(s) that get upregulated do not necessarily depend on the degree of sequence homology or mutated gene length in a linear fashion. Future studies may explore whether alternative paralog- or knockout-target-specific characteristics are associated with upregulation after mutation.

### Building a minimal network model of the effects of nonsense-induced transcriptional compensation

We demonstrated the existence of transcriptional adaptation in mice and humans across multiple contexts. While such publicly-available datasets provide an important initial view of nonsense-induced transcriptional compensation, several questions related to the phenomenon’s downstream effects, including the penetrance of phenotypes, remain unanswered. For example, under what conditions is transcriptional adaptation capable of inducing robustness across a population of cells, in that the compensating paralog expression precisely mimics wild-type expression at a single-cell level? Robust paralog expression may not be sufficient, as paralog activity (e.g., that of a paralogous enzyme or a transcription factor) can differ substantially from the original gene. Similarly, gene regulatory network effects can result in distributions of effector molecules in single cells that are non-trivial to predict, yet can have profound phenotypic implications. For example, mutations in regulators of *C. elegans* intestinal fate can result in downstream effector expression heterogeneity, further dependent on the continued function of other regulatory network components ^53^. Therefore, it is important to identify the major control knobs that may confer robustness, or lack thereof, to obtain a mechanistic understanding of single-cell variability underpinning transcriptional adaptation. To address this gap, we built a theoretical framework to model the ensemble of single-cell adaptation possibilities with a minimal set of stochastic biochemical reactions.

We chose to model cells in which a gene that exhibits nonsense-induced transcriptional compensation controls the expression of a downstream effector molecule. In our initial minimalistic model (Methods), we simulate transcription of an upstream regulator, A, with a paralog, A’, exhibiting nonsense-induced transcriptional compensation, and a downstream target, B, in a diploid genome (Figure 3A). Gene product A in wild-type regulates the transcription of downstream pathway member, B. Mutation of A is compensated for by nonsense-mediated expression enhancement of A’, which also regulates transcription of B when present (Figure 3A). To model the effect of nonsense-mediated expression enhancement of A’ on B, we used an expanded version of the telegraph model of transcription ^54^ as a building block in our model: each gene can reversibly switch between a transcriptionally inactive state (to which, r_off_) and one or more active states (to which, r_on_) (Figure S4A). When active, the gene product is transcribed in a Poisson process at a rate (r_prod_). Degradation of each product also occurs as a Poisson process (r_deg_). We specify the directed interaction between mutated A regulating the transcription of paralog gene A’, which represents nonsense-induced transcriptional compensation, by adding a parameter (r_add_ ^NITC^) with dependency on the real-time abundance of mutated A gene product modified by a Hill function (Hill coefficient n), to account for the nonlinearity of gene regulatory interactions. We combine steps leading to transcription by making the quasi-equilibrium assumption, commonly used in models of gene regulatory networks due to differences in individual reaction timescales ^55,56^. We represent the differential regulation of B by A and A’ by specifying two distinct active states for B: the active state directed by A and the active state directed by A’. The active states of B each have a respective production rate. In sum, our minimalistic model includes 12 varying parameters and 1 fixed parameter. We condensed the parameter search space to 8 independent and interpretable variables by focusing on parameter relationships in relation to a subset of critical network parameters (see Methods).

**Figure 3:**
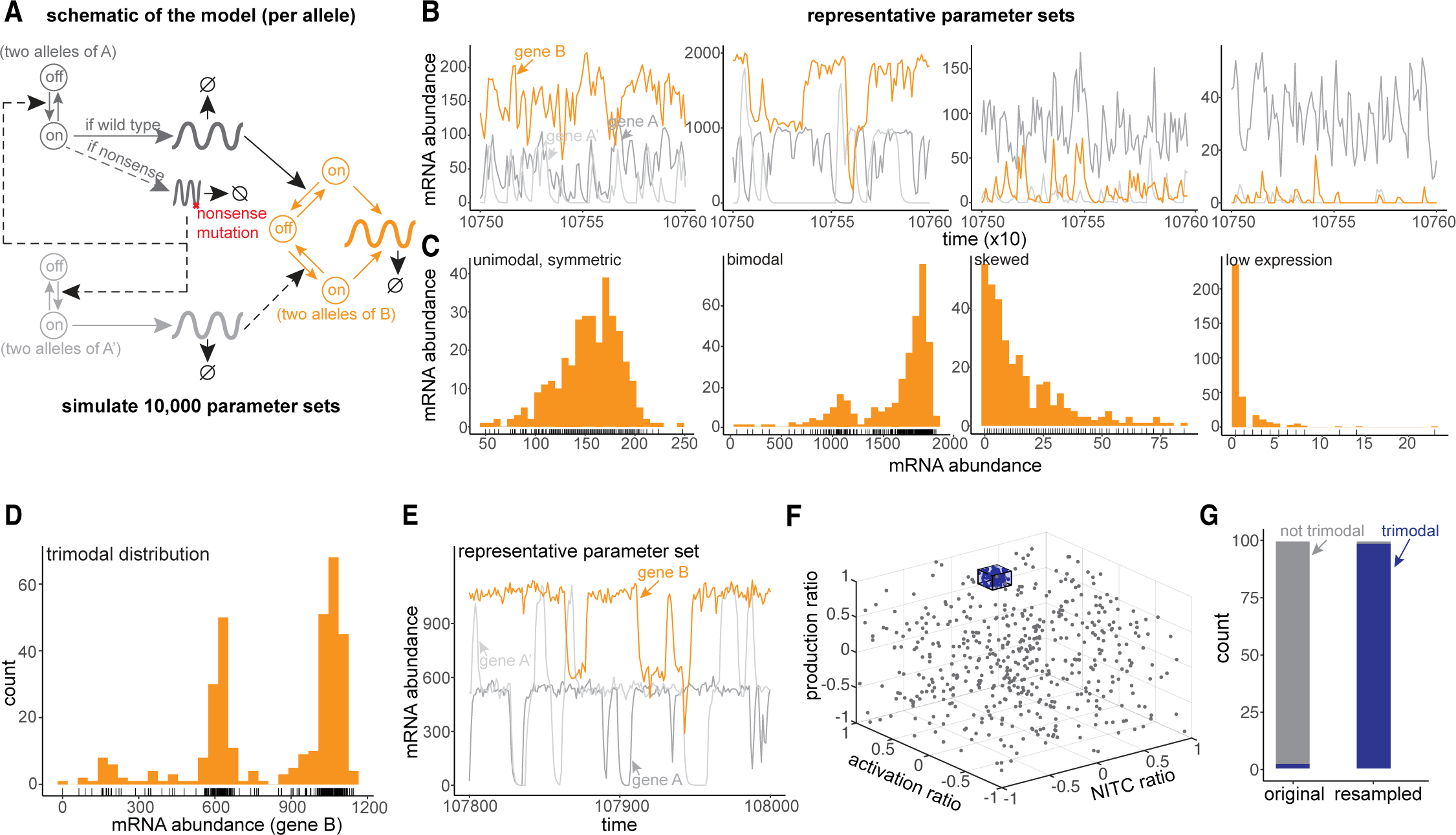
Outputs of simulated gene regulatory networks with transcriptional adaptation and enrichment of distribution shapes in parameter subspaces A. Schematic of a gene regulatory network with transcriptional adaptation to mutation. Two alleles of each gene, with bursty transcription of gene products at each allele. A mutated reference gene, A (dark gray), regulates downstream effector gene B (orange). When mutated, nonsense copies of A product upregulate a paralog of A, called A’ (light gray). A’ can also regulate B, albeit with different strengths. Hill functions are used in propensities for regulatory relationships between gene products and target alleles. See Methods for full model specification. Parameter descriptions in the table at right of the panel. B. Example simulation output and inference of single-cell expression distributions from pseudo-single-cells taken every 300 time-steps. See Methods. C. Example classification of gene expression distribution shapes. See Methods for classification algorithm. D. Example trimodal gene expression distribution. E. Example simulated gene expression tracing for trimodal distribution parameter set. F. Parameter subspace around the initially identified trimodal distribution’s parameter set. G. Enrichment of trimodal gene expression distribution shapes after sampling parameter sets from the subspace versus from the full parameter space.

### Gene expression distribution shape varies widely across the parameter search space

To understand how relationships between the parameters in the model relate to network output, we scanned the following independent variables: the basal on-rate of A, the production rate when active (applied to all genes), the Hill coefficient n (applied to all genes), the r_add_ of A on transcription of B, and four parameter ratios of interest (Figure S4A). Four independent parameter ratios were: the ratio of r_add,NITC_ to the on-rate of A (how strong is NITC relative to basal regulation of A), the ratio of the on-rate of A to the off-rate (how rapidly does A’s expression burst cycle), the ratio of r_add_ of A on B to r_add_ of A’ on B (what are the relative strengths of A and A’ causing B burst activation), and the ratio of r_prod_ of B in the A-directed on state to the r_prod_ of B in the A’-directed on state (the relative activities of B on states) (Figure S4A). We used the Gillespie algorithm to numerically simulate network output under different parameter values ^57^.

We chose parameter search ranges based on several studies empirically documenting transcriptional burst kinetics, when available ^54,58–60^. A recent study of transcriptome-wide allele-specific expression in single cells inferred telegraph model parameters relative to transcript degradation rates ^58^. We therefore fixed the degradation rate at 1, concordant with Larsson and colleagues’ analysis, and picked ranges of the burst-on rate of A, the on-off rate ratio, and production rate based on the range of parameters inferred by Larsson and colleagues ^58^ (Figure S4B-D). Other parameter ratios were similarly picked to span two orders of magnitude while also ensuring at least partial concordance with the parameter ranges observed for simple telegraph models ^54,58–60^.

We simulated our stochastic transcriptional model to capture and quantify the emergent single-cell variability within the minimal gene regulatory networks (Figure 3B). Since the simulations are expected to be ergodic, we reasoned that we could condense long-timescale traces into “single-cell-like sub-simulations” that are independent of each other (Figure S5A-C, Methods) ^54^. Indeed, we found no significant autocorrelation in the single-cell-like sub-simulations, which, in turn, enabled us to get population-level distributions of counts of each gene per parameter set. Briefly, with static snapshots taken every 300 timesteps, we simulated hundreds of single-cell-like gene expression profiles in wildtype, heterozygous mutant, and homozygous mutant genotypes. We summarized the expression levels per gene per genotype in each simulation with manually annotated empirical distribution classes (Figure 3C). We confirmed that average gene expression levels matched steady state approximations for the corresponding system of differential equations using the ode45 solver in MATLAB (Figure S6A-C).

Next, since our simulations resulted in several tens of thousands of distributions for each of the gene products, we needed an automated method for describing expression distributions. First, we used summary statistics to describe the moments and other features of expression distributions including mean, coefficient of variation, skewness, bimodality coefficient (a statistic correlated with multimodality), gini coefficient, and entropy. For expressed genes, the values of statistics varied widely across parameter sets (Figure S7). Manual inspection of random samples showed that the distributions tended to fall into 5 general classes of distribution shape: low-expression, unimodal symmetric, left-skewed, right-skewed, and bimodal (or multimodal) (Figure 3C). In order to systematically classify distribution shapes, we developed a heuristic algorithm based on summary statistics and verified the accuracy with manual checks (80-100% per class, see Methods, Figure S8A). We explored distribution shapes and their respective summary statistics to arrive at classification rules predicting distribution shape. The rules included mean, skewness, bimodality coefficient, and bimodality coefficient normalized to mean expression level using LOESS (since there was systematic variation in bimodality coefficient with the mean) (see Methods). We then asked if we could use this classifier to capture the frequency of distribution shape change after mutation as a proxy for whether there was robustness in distribution shape. For gene product B, we found that while many simulations demonstrated shape class changes after mutation, a large number instead did not (Figure S4E). This robustness in distribution shape after mutation could translate into downstream phenotypic robustness.

In our initial set of simulations, we considered paralogs with no basal expression, i.e., paralogs regulated exclusively by transcriptional compensation. We wanted to check if paralogs with some basal expression, like many observed paralogs in real world datasets, display similar diversity in distribution shapes and robustness of distribution shape to mutation ^61^. We ran 10,000 more simulations, keeping the original 8 independently varying parameters and ratios in their original ranges. We added a 9th independent variable: the ratio of the basal burst-on rate of A’ relative to that of A. The second set of simulations produced all 5 distribution classes observed in the first set.

Notably, more parameter sets preserve the distribution shape of the downstream target, B, after mutation, than in the first set of simulations (Figure S9). A larger fraction of robustness-to-mutation parameter sets suggests that having non-zero basal paralog expression, representative of gene A-A’ redundancy, may help ensure robustness to mutation.

### Gene expression distribution shape depends on model parameters

The shape and variability of a single cell’s gene expression distribution can affect phenotypic penetrance and disease progression. Therefore, we are most interested in understanding what influences gene expression distributions in a mutant, heterozygous, state. We questioned how compensatory intrinsic gene regulatory mechanisms might control the shape of emerging population-level distributions. As we observed a diversity of gene expression distribution shapes, we checked whether there were any associations between independent model variables (i.e., core parameters and parameter ratios) and gene expression distribution shapes. Since gene B represents the model network output and the heterozygous mutant state is the genotype of primary interest in this study, we focused on summary statistics describing distribution shapes for gene B in the heterozygous genotype. Of the summary statistics, we first considered bimodality coefficient since it is correlated with whether a distribution has more than one mode. We were particularly interested in scenarios in which a predictable, unimodal population can retain its unimodality after mutation (and therefore has a low bimodality coefficient). We found that the ratio of B production rates in A- versus A’-directed on-states was more strongly correlated with bimodality coefficient than other relevant variables (Figure S8B). Indeed, the absolute value of the log ratio in our model parameterization (see Methods), was correlated with bimodality coefficient (r = 0.32).

Beyond associations between individual model parameters and distribution summary statistics, we wondered whether combinations of parameters or parameter ratios were more likely to give rise to particular distribution shape classes upon mutation. We chose to focus on unimodal symmetric, as this distribution shape is most reflective of a homogeneous, unskewed population, with non-zero expression. Therefore, we trained a decision tree classifier on the sampled parameter sets (see Methods) to identify the model parameter and ratio combinations most predictive of whether gene B in the heterozygous genotype would be unimodal symmetric class (see Methods). We found a total of 31 significant decision rules up to 6 layers deep per combination, resulting in 33 groupings of parameter sets (Figure S10). All included independent parameters and ratios that were present in at least one decision rule. The combination with the highest fraction of unimodal symmetric gene B distributions in the heterozygous state had 6 decision rules, including requiring sufficiently strong r_add,NITC_ (i.e., a low ratio of r_on,basal_ of A to r_add,NITC_ acting on A’). Consistent with bimodality coefficient correlation results, other subspaces enriched for unimodal symmetric shapes included rules restricting the ratio of A- and A’-directed B production rates to be closer to 1.

In further exploring the data, we identified an unexpected gene expression shape in one out of more than a hundred randomly inspected gene B expression distributions: trimodal with 3 non-zero modes (Figure 3D,E). Trimodality was unexpected because *a priori* the expected possible regulation regimes for gene B are A-driven, A’-driven, and neither-driven, with the last having no basal expression.

Examination of the gene expression traces suggested that the rate of B mRNA degradation was too slow to reach zero before an allele was activated by A or A’ (Figure 3E). This result raises the plausibility of the existence of a triad of phenotypic outcomes, and underscores the emergent complexity inherent to even relatively simple gene regulatory networks. To verify whether this distribution was associated with the specific parameter combination, i.e., subspace within parameter space for this parameter set, we sampled directly around the parameter set with defined ranges smaller than the original search space ranges (Figure 3F, see Methods). Indeed, we found that essentially all (98%) parameter sets sampled within the narrow subspace ranges displayed similar trimodality, while a new random sample of the full original parameter space resulted in only 1% of sampled simulations displaying trimodality of B (Figure 3G). This analysis further established that we can use parameter set combinations to predict distribution shapes.

### Gene expression distribution robustness to mutation is dependent on model parameters

We next asked whether we could identify model predictors of robustness to mutation, insofar as robustness could exist in our model (Figure 4A). We performed decision tree analysis again, but this time restricted to parameter sets in which gene B was unimodal symmetric in the wild-type state. We trained a classifier on whether or not a given parameter set resulted in gene B remaining unimodal symmetric in the heterozygous mutant genotype. This criterion would ensure that the downstream gene product retained its distribution shape even after the upstream regulator was mutated. We found a limited number of significant decision rules for unimodal symmetric robustness: a total of 11 decision rules, up to 5 layers deep, with 12 total groupings (nodes) of parameter sets (Figure 4B).

**Figure 4:**
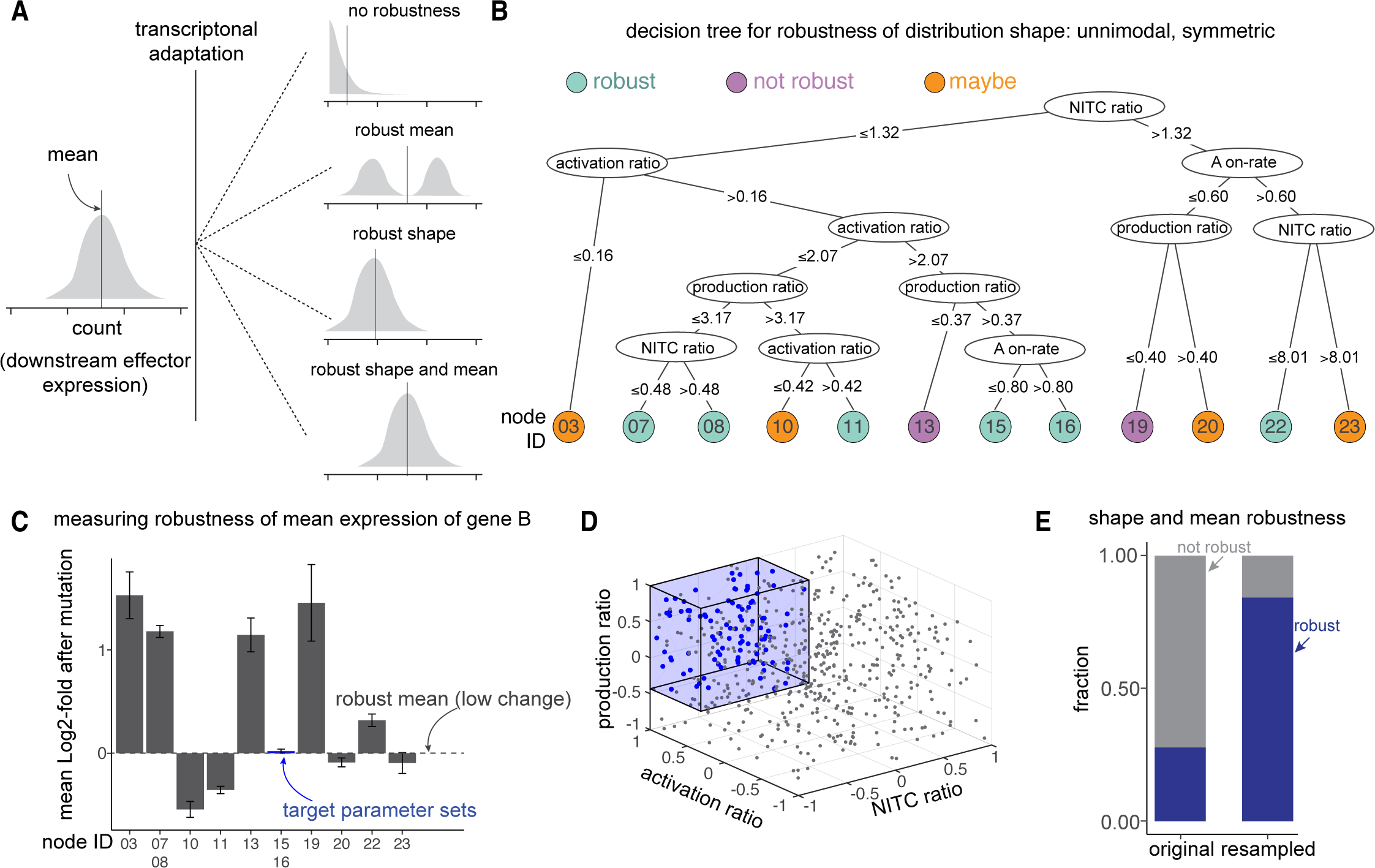
Robustness to mutation dependency on model parameters A. Analysis schematic: under what network conditions does the network output (i.e., distribution of B expression levels) remain unchanged, either in shape or in average expression level, after mutation of gene A? B. Decision tree trained on model parameters to classify parameter sets, restricted to those in which B is unimodal symmetric in the wildtype genotype, by whether B distribution remains unimodal symmetric or if it changes distribution shape class. Nodes marked “robust”: >70% of considered parameter sets robust, “maybe”: 30-70% robust, “not robust”: <30% robust. C. Mean B expression changes after mutation in each decision tree leaf for unimodal symmetric robust parameter sets. D. Parameter subspace bounded by decision tree rules in (C), resampled. E. Enrichment of robustness of both shape and mean for unimodal symmetric distributions of B after mutation, subsampled from the full parameter space and from the subspace marked in (D). Robustness of mean here defined as absolute log_2_ fold-change after mutation < 0.35.

Next, we posited that besides the distribution shape, average expression of B must be accounted for when measuring robustness. We found that among 4 nodes with shape robustness to mutation, only nodes 15 and 16 demonstrated preservation of average expression level after mutation. Therefore, we identified exactly one parameter subspace that, for wild-type unimodal symmetric parameter sets within it, demonstrated robustness of distribution shape and average expression level. This parameter space is bounded by the three condensed decision rules: 1) ratio of r_add_ ^NITC^ to r <= 1.32 (up to moderate strength NITC), 2) ratio of r_add_ of A on B to r_add_ of A’ on B > 2.07 (relatively less frequent A’-directed bursting of B), and 3) ratio of r_prod_ of B in the A-directed on state to the r_prod_ of B in the A’-directed on state >0.37 (A’-directed B production rate can be either weaker or stronger than A-directed, but not by too much) (Figure 4C). To confirm these results, we generated new parameter sets within the node 15+16 parameter subspace (Figure 4D). We found that the newly sampled parameters were strongly enriched for robustness of shape and average expression level as compared to all parameters (Figure 4E).

### Qualitatively similar rules hold for distributions and robustness across multiple biological contexts

Many human and mouse genes have multiple annotated paralogs that could in principle contribute to transcriptional adaptation upon mutation ^40^. We therefore wondered whether an expanded gene regulatory network model, with multiple paralogs of the ancestral regulator A, would demonstrate similar diversity of resulting expression distribution shapes and robustness to mutation of A (see Methods, Figure S11A). Indeed, we observed a diversity of distribution shapes across the full parameter space, with all 5 previously observed classes represented (Figure S11B,C), and robustness results were qualitatively concordant with single paralog network simulations (Figure S12, S13).

Many gene regulatory networks also include transcription factors with repressive effects on downstream targets ^62^. We wondered whether transcriptional adaptation could be important for preserving network output after mutation of a repressor (see Methods; Figure S14A). Here again, we observed similarly diverse distribution shapes across the full parameter space (except left-skewed distributions) (Figure S14B,C), and a subset of parameters (albeit a narrower set than activator network) enabled robustness (Figure S15,16). Further work may elucidate regulatory constraint differences between activator- and repressor-based networks with compensation.

### Robustness of regulons at bulk and single-cell resolution for transcription factors exhibiting transcriptional adaptation

We wondered whether the gene expression robustness observed in simulations of gene regulatory networks with transcriptional adaptation was observable in experimental data. Specifically, we wanted to know if there was robustness of expression distributions for downstream targets of transcription factors—which can activate or repress downstream genes—that showed signs of transcriptional adaptation via paralog upregulation (Figure 5A). To address this question, we isolated transcription factors from the bulk RNA-seq and Perturb-seq single-cell RNA-seq datasets previously analyzed for paralog upregulation (Table S1 and ^45^).

**Figure 5:**
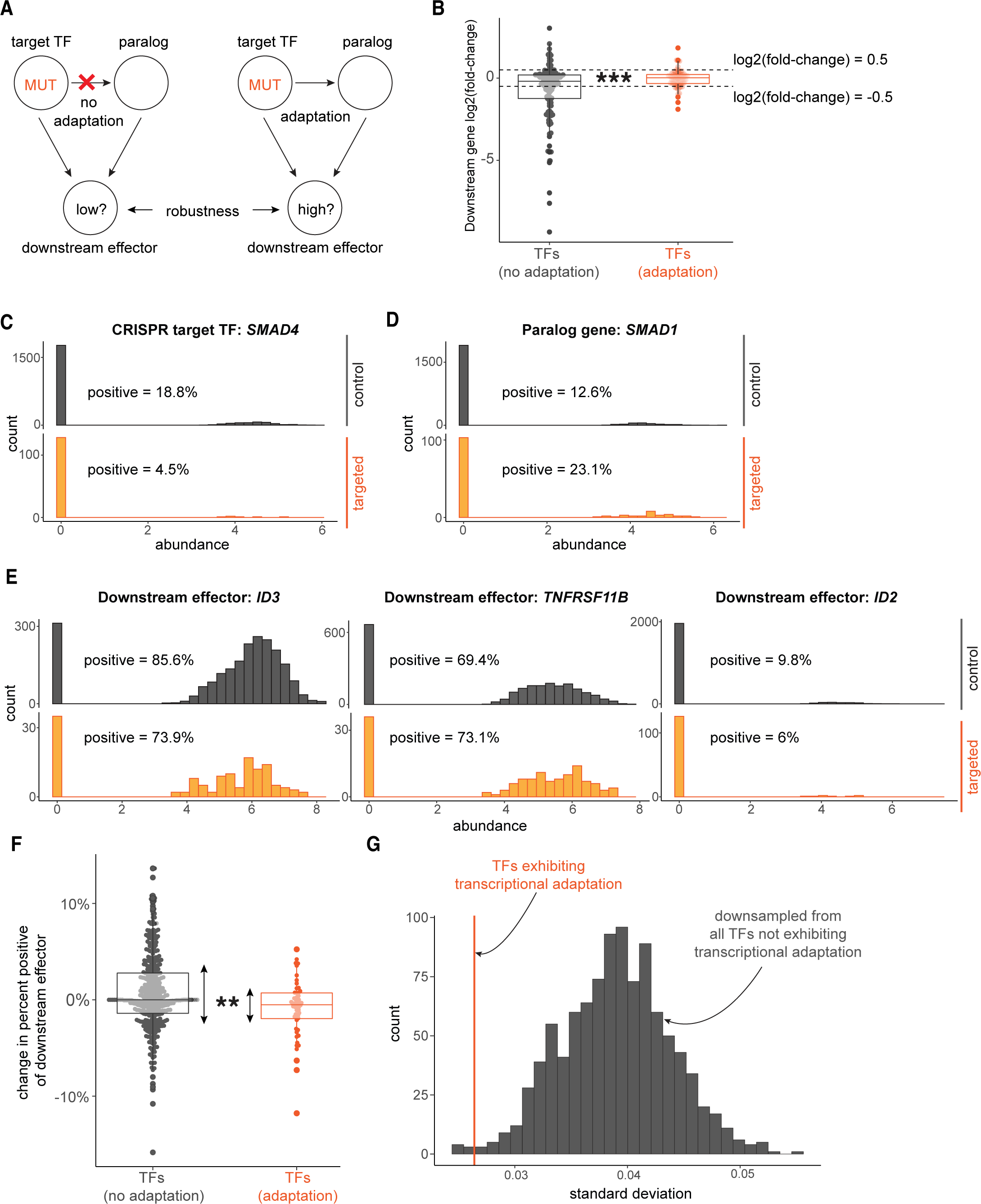
Robustness of regulon expression associated with transcription factor transcriptional adaptation A. Analysis schematic: is there different regulon gene expression robustness after mutation of upstream transcription factors, associated with whether the transcription factors demonstrate transcriptional adaptation by paralogs? B. Change in gene expression for each regulon gene after reference CRISPR transcription factor target mutation, compared against respective controls (log2(fold change) from DESeq2). Each point represents one downstream regulon member gene. Gray: regulon genes downstream of CRISPR target-paralog pairs not appearing to be involved in transcriptional adaptation. Red: regulon genes downstream of CRISPR target-paralog pairs with apparent transcriptional adaptation. Regulon genes called differentially expressed if DESeq2 adjusted p-value < 0.05 and abs(log2(fold change)) > 0.5. Fisher’s exact test for difference of odds of differential expression between the two groups of CRISPR target regulons, p = 1.82 x 10^-4^. C. Gene expression distributions of a representative transcription factor that demonstrates possible transcriptional adaptation: *SMAD4*, in non-template controls (“control”, gray) and in *SMAD4*-guide-treated cells (“targeted”, orange). D. Gene expression distributions of a representative top-100 paralog gene of a transcription factor that demonstrates possible transcriptional adaptation: *SMAD1*, a paralog of *SMAD4*, in non-template controls (gray) and in *SMAD4*-guide-treated cells (orange). E. Gene expression distributions of three representative regulon genes of a transcription factor that demonstrates possible transcriptional adaptation: *ID3, TNFRSF11B, and ID2,* regulon genes of both *SMAD4* and *SMAD1*, in non-template controls (gray) and in *SMAD4*-guide-treated cells (orange). F. Change in gene expression for each regulon gene after reference CRISPR transcription factor target mutation, compared against non-template controls (i.e., percent-positive in CRISPR targeted cells minus percent-positive in non-template control-treated cells, for genes 75% or less in non-template controls). Each point represents one downstream regulon member gene. Gray: regulon genes downstream of CRISPR target-paralog pairs not appearing to be involved in transcriptional adaptation. Red: regulon genes downstream of CRISPR target-paralog pairs with apparent transcriptional adaptation. Asymptotic test of difference in coefficient of variation for groups of unequal size, p = 0.0017. G. Empirical distribution of standard deviation of change in percent-positive for downsampled (n=49, same as transcriptional-adaptation group size) no-transcriptional-adaptation group regulon members without replacement, 1,000 downsamples, in gray. Observed standard deviation of change in percent-positive transcriptional-adaptation group standard deviation to this distribution, in red.

For the bulk RNA-seq datasets, we first checked whether any CRISPR targets with significant paralog upregulation were transcription factors. Next, we searched the DoRothEA regulon database for common downstream targets of CRISPR targets and their paralogs (i.e., common regulons) ^63,64^. We found common regulons with high-quality annotations for 4 target-paralog pairs for targets demonstrating possible transcriptional adaptation: *RUNX3-RUNX1, SP1-SP4, ZEB2-ZEB1,* and *Myc-Mycn*. In parallel, we repeated regulon searches for all target-paralog pairs for targets that did not appear to demonstrate transcriptional adaptation, as well. We hypothesized that transcriptional adaptation by transcription factor paralogs should, on average, buffer extreme changes in the expression of downstream genes after CRISPR target mutation. To test our hypothesis, we calculated differential expression of common regulon genes after CRISPR target knockout. We found that common regulon genes downstream of transcription factors with signs of transcriptional adaptation were significantly less often differentially expressed (10.2% of regulon genes) than those downstream of transcription factors without signs of transcriptional adaptation (34.1% of regulon genes; Fisher’s Exact Test p-value = 1.82 x 10^-4^; Figure 5B).

For the Perturb-seq dataset, we asked whether any of the top-100 upregulated paralogs were paralogs of known human transcription factors. Six pairs of top-100 CRISPR targets-paralog genes spanning 4 distinct CRISPR targets were transcription factors. We then searched the DoRothEA regulon database for common downstream targets of CRISPR targets and their paralogs (i.e., common regulons) ^63,64^. We found overlapping regulons with high-quality annotations for 3 of the 6 top-100 target-paralog pairs: *IRF4-IRF1*, *SMAD4-SMAD1*, and *TFAP2A-TFAP2C* (Figure 5C-E). For each regulon, we then plotted single-cell transcript abundances for the CRISPR target (Figure 5C), the paralog (Figure 5D), and downstream genes (Figure 5E) both in non-template control cells and in cells treated with guides specific to the CRISPR target. By inspection, we observed that some downstream genes demonstrated fully robust, consistent gene expression distribution shapes, while others demonstrated partial expression robustness with possible new modes of gene expression (i.e., the possible presence of bimodality or multimodality), or lower expression overall (Figure 5E).

As with the bulk RNA-seq datasets, we hypothesized that transcriptional adaptation by transcription factor paralogs would, on average, buffer extreme changes in downstream genes after CRISPR target mutation. Therefore, we repeated regulon searches and calculated the change in downstream gene expression for target-paralog pairs that did not appear to demonstrate transcriptional adaptation and for those exhibiting transcriptional adaptation. Compared to regulons of CRISPR target transcription factor-paralog pairs that do not appear to demonstrate transcriptional adaptation, the spread of gene expression changes downstream of top-100 target-paralog pairs was indeed narrower (p = 0.0017, see Methods) (Figure 5F,G). In summary, transcriptional adaptation by paralogs of mutated transcription factors is associated with buffering of extreme expression changes in their mutual downstream regulon genes, in both bulk and single-cell datasets.

## Discussion

We developed a computational framework integrating bioinformatic analysis, mathematical modeling, and machine learning to uncover the genome-wide prevalence and gene regulatory constraints on a recently reported kind of transcriptional adaptation, nonsense-induced transcriptional compensation, in single mammalian cells. We found transcriptional upregulation of paralogs after reference gene mutation to be pervasive, but not necessarily ubiquitous, across cell types and contexts, including cancer, development, and cellular reprogramming. Furthermore, the genes identified as exhibiting possible transcriptional adaptation were neither associated with any single signaling pathway nor did they exhibit any observable molecular functional congruence between each other. Our relatively parsimonious model consisting of transcriptional bursting and stochastic interactions between genes in a biallelic compensatory network could produce a range of population-level distributions of downstream targets upon compensation, underscoring the complex ensemble of fate-space that compensatory networks can access. Finally, our regulon robustness results synthesize two separate earlier analyses: paralog upregulation bioinformatic analysis and single-cell network simulations, in that transcriptional adaptation was associated with downstream regulon gene robustness after transcription factor mutation. Collectively, our computational framework provides a basis for further mechanistic experimental and computational studies on the origins and manifestations of nonsense-mediated transcriptional adaptation.

Recent advances in sequencing and genome editing technologies have enabled perturbation and profiling of the molecular makeup of single cells at unprecedented throughput. For greater resolution of differences between genetic perturbation methods, high-throughput parallel treatments of the same target genes with RNA interference, CRISPRi, and Cas9-based knockouts could reveal specific effects of post-transcriptional, epigenetic, or mutation-based methods. Perhaps such parallel experiments could reveal transcriptional adaptation, or yet unknown mechanisms, by which cells retain robustness to genetic perturbations. Similarly, the adoption of recent single-cell CRISPR screening frameworks, such as CROP-seq and Perturb-seq ^42–44^, coupled with high-depth sequencing can lead the way in identifying single-cell manifestations of transcriptional adaptation. These experimental findings can, in principle and if at higher quantitative resolution, be projected onto the distributions from our theoretical formulations. Given the non-linearities associated with sequencing datasets, *bona fide* gene targets identified from sequencing studies can be tested in single cells with single-molecule fluorescent *in situ* techniques, which measure the absolute expression counts in individual cells for greater quantitative resolution ^65^.

The mapping between simulation and wet-lab experiment can uncover plausible network and parameter constraints for individual compensating genes. Such mappings could also help explain incomplete phenotypic penetrance reported in association with transcriptional adaptation. Another set of questions center around whether gene length, number of introns and exons, chromosomal locations, and chromatin landscape play a role in which gene families exhibit nonsense-induced transcriptional compensation. Furthermore, such mappings can help with the design and interpretation of functional genetic screens by taking into account genes known to be exhibiting transcriptional adaptation and the extent of its impact. The breadth of genes that appear to have transcriptional compensation also invites study of potential negative consequences of nonsense-induced paralog—or other related gene—upregulation. Might some compensatory changes be deleterious, and if so, could such deleterious changes explain select negative phenotypes previously ascribed to haploinsufficiency or gene dosage effects ^66^? In a similar vein, our framework could be extended to analyzing cases where paralogs are downregulated upon Cas9-induced nonsense mutations, potentially revealing new biology.

One limitation of our work is that a majority of the analysis was performed on datasets from bulk RNA sequencing studies, limiting a quantitative single-cell mapping with simulations. As single-cell sequencing datasets, and single-cell transcriptomics via other methods that enable absolute expression counts, such as SeqFISH, MERFISH, and optical pooled screens ^67–69^, become more accessible, bioinformatic analysis can inform model architecture and parameters and move towards more predictive models. Another limitation of our framework is that we focused primarily on mice and human datasets given the breadth of available datasets. In principle, our bioinformatic pipeline can be generalized to include other animal systems to reveal both species-specific and universal gene targets displaying transcriptional compensation ^14,15^. Lastly, simulations of gene regulatory networks are inherently simplifying, and while we specified reactions and assumptions that have been shown to model small numbers of interacting genes well ^59,60,70^, these models do not account for all regulatory interactions in a cell explicitly ^32,54,71^.

## Acknowledgements

We thank members of the Goyal lab for insightful discussions related to this work. We also thank Lea Schuh (Helmholtz Munich), Karun Kiani (University of Pennsylvania), and Granton Jindal (UCSD) for discussions and manuscript comments. We thank Aviv Regev (Genentech) for pointing us to relevant datasets. YG acknowledges support from Northwestern University’s startup funds and the Burroughs Wellcome Fund Career Awards at the Scientific Interface. NB acknowledges support from NIH T32 GM144295, T32 GM142604, and funding to YG. We thank all authors who published accessible transcriptomics datasets.

## Author Contributions

YG and IAM conceived and designed the project. IAM designed, performed, and analyzed the simulations under the guidance of YG. IAM conceptualized the analysis of publicly available datasets, and NB identified datasets and performed the analysis under the guidance of IAM and YG. MEM performed a subset of analysis and draft review with input from IAM and YG. IAM and YG made the figures. YG and IAM wrote the paper with input from MEM and NB.

## Methods

### Selection of CRISPR-Cas9 transcriptomics datasets

We searched published literature, preprints, and the Gene Expression Omnibus (GEO: https://www.ncbi.nlm.nih.gov/geo/) for RNA sequencing datasets generated from experiments designed to measure differential gene expression after Cas9-mediated knockout of a gene of interest. We used the following search terms: “CRISPR/Cas9” or “CRISPR-Cas9” and “RNA-seq”. We focused our search for publicly available data on the GEO RNA-seq Experiments Interactive Navigator (GREIN). GREIN contains processed data from thousands of GEO entries with human, mouse, and rat RNA-seq samples using a common pipeline for alignment, quality control, and transcript quantification ^72^. In GREIN we used search terms “CRISPR/Cas9” or “CRISPR-Cas9”. Based on these search results, we manually checked the experimental designs of more than 200 publicly available RNA-seq experiments. We only considered experiments in which 1) there was CRISPR/Cas9-based knockout of the target, in which the stated strategy was not to target intergenic regulatory sequences, 2) annotated matched control samples treated with non-template control gRNA, and 3) multiple replicates of both targeted and control RNA-seq samples. We prioritized studies with multiple knockout targets, which enabled us to check for the presence or absence of paralog upregulation for as many targets as possible. For the few non-GREIN datasets included here, we only considered studies that provided mapped read counts per gene ID or, optimally, also provided processed differential gene expression calculations. Ultimately we found 36 studies with datasets meeting inclusion criteria described above, described in Table S1 ^14,41,43,52,73–100^.

### Identification of paralogs of knockout targets

For bulk RNA-seq datasets, we queried the Ensembl database version 110 for paralogs of knockout targets, using Ensembl REST API (Version 15.6) ^40^. We searched by Gene Symbol, and extracted paralogs of all returned Ensembl IDs. We extracted both gene identifiers and Ensembl-annotated coding sequence overlap percentages between knockout targets and each of their paralogs. Ensembl gene IDs were then converted to standardized gene names using g:Profiler (Version e109_eg56_p17_1d3191d) ^101^. For Perturb-seq data from ^45^, we used the BiomaRt package v2.40.5 in R v3.6.1 to search Ensembl version 105 for paralogs, searching by Gene Symbol and extracting all returned paralog Gene Symbols ^102^.

### Differential gene expression assessment

We wanted to identify differentially expressed genes across the dozens of knockout samples we reanalyzed. We used DESeq2 for differential expression analysis ^103^. When available, we used author-provided gene expression change calculations based on DESeq2 (for results from ^52^). For all remaining datasets, for which DESeq2 results were not already available, the authors did provide mapped count data on GEO and/or they were available on GREIN. For these count-based results we implemented DESeq2 ourselves, using the PyDESeq2 package, using default settings, comparing knockout samples against the matched controls from their respective studies ^104^.

For these studies, we implemented filters to consider knockout targets that we would *a priori* expect to have some detectable loss of gene dosage that would need to be compensated for by transcriptional adaptation. We first confirmed that the average library size of considered samples was at least approximately 1 million reads per sample. We then included genes only if they were expressed at a level of 10 raw counts or higher across all samples. We chose to classify paralogs as upregulated if DESeq2 reported an adjusted p-value <=0.05 and a log_2_ fold-change >= 0.5. In supplementary analyses we also show results when paralogs are classified as upregulated using either 1) only the adjusted p-value <=0.05 filter, or 2) adjusted p-value <=0.05, log_2_ fold-change >= 0.5, and basemean >= 10 filters. The final analysis included all knockout target genes with any significant paralog differential expression, up or down, irrespective of log_2_ fold-change.

### Estimation of expected frequency of paralog upregulation per knockout target

The null hypothesis for analysis of paralog upregulation is that for any group of paralogs of a knockout target, the number of paralog genes upregulated after knockout is simply reflective of randomly selecting any similarly expressed genes in the dataset and checking whether they were upregulated. In order to check whether the paralog upregulation pattern observed for a particular knockout target was reflective of randomly selecting similarly expressed genes in the dataset (instead of the paralogs), we developed an algorithm for bootstrapping the null distribution of paralog upregulation frequency for each knockout target. For each knockout target, the algorithm is implemented as follows:

1. Rank order all genes in the dataset by basemean across all samples
2. For each paralog, randomly select a gene within 50 ranks
3. For each randomly selected similarly expressed gene, check the fold change after knockout and whether the gene qualifies as upregulated based on dataset-specific thresholds (in figure legends)
4. Count the number of upregulated randomly selected genes and divide by the total number of paralogs for a bootstrap sample of the paralog upregulation frequency
5. Repeat steps 2-4 10000 times to build an empirical null distribution of the paralog upregulation frequency
6. To calculate a p-value, calculate the fraction of the empirical null distribution that is at least as large as the observed fraction of paralogs that are upregulated

### Perturb-seq-based single-cell gene expression reanalysis

We wanted to identify possible changes in single-cell gene expression distributions after knockout of a library of CRISPR targets. Therefore, we reanalyzed Cas9-based pooled knockout single-cell RNA-seq, Perturb-seq, data from ^45^. We downloaded published processed log-transformed UMI-based transcript quantification tables from https://singlecell.broadinstitute.org/, accession SCP1064, “Control” condition, last accessed June 15, 2023. For the main analysis, we only considered knockout targets with sufficient cells for a minimal quantitative analysis: a minimum of 2000 UMI per cell, at least 30 cells total, with no fewer than 5 cells annotated in any one of the three included targeting guide RNAs. In a supplementary analysis to identify lower-confidence targets with possible transcriptional adaptation, we considered removing the 30-cell and 5-cell filters. For transcript quantification, within each guide, we either counted the number of cells with non-zero expression and divided by total cells for that guide (for percent-positive), or we averaged expression levels over all cells for that guide (for mean). Within each gene within a given condition (nontemplate controls or targeted for a given gene), we averaged over all appropriate guides.

### Regulon robustness analysis

For bulk RNA-seq data-derived regulon gene expression analyses, we focused on human and mouse transcription factors, as defined by the most recent version of AnimalTFDB3, last accessed November 21, 2023 ^105^. We searched for overlapping regulons between a knockout target and the paralog gene of interest in DoRothEA, only considering downstream genes with annotation confidence level A, B, or C (out of a possible range of A-E, see original source for evidence level descriptions) ^63,64^. We compared regulon genes for transcription factor CRISPR target-paralog pairs in the bulk RNA-seq dataset that demonstrated possible transcriptional adaptation as defined by significant paralog upregulation frequency (p<0.1, see Estimation of expected frequency section, above). There were 68 annotated regulon genes across the four target-paralog pairs with transcriptional adaptation versus 138 annotated regulon genes for all target-paralog pairs without transcriptional adaptation. Regulon genes were considered differentially expressed if DESeq2 adjusted p-value < 0.05 and abs(log2FoldChange) > 0.5. We calculated a p-value between the groups of regulon genes using Fisher’s exact test, testing whether the odds of regulon gene differential expression were different between the transcriptional-adaptation and no-transcriptional-adaptation groups.

For Perturb-seq data-derived gene expression distribution analyses, we chose to focus on human transcription factor genes, as defined by the most recent version of AnimalTFDB3, last accessed July 28, 2023 ^105^. We searched for overlapping regulons between a knockout target and the paralog gene of interest in DoRothEA, only considering downstream genes with annotation confidence level A, B, or C (out of a possible range of A-E, see original source for evidence level descriptions) ^63,64^. We compared regulon genes for transcription factor CRISPR target-paralog pairs in the Perturb-seq dataset that demonstrated possible transcriptional adaptation as defined by having a top-100 paralog, versus all pairs that that did not demonstrate transcriptional adaptation, as defined by both not having any top-100 paralogs and being a pair in the interquartile range of changes in paralog expression after knockout. There were 55 annotated regulon genes for the three target-paralog pairs with transcriptional adaptation versus 439 annotated regulon genes for all target-paralog pairs without transcriptional adaptation.

The average difference in expression of regulon genes in both groups was approximately zero, with a spread of values about that mean (Figure 5C). In order to test for transcriptional adaptation-associated buffering of gene expression changes of regulon genes after CRISPR target mutation, we performed two checks for significance of a difference in the spread size between the two groups; a larger spread would indicate a larger number of extreme expression changes. The first test was an asymptotic test of difference in coefficient of variation for groups of unequal size, using the cvequality v0.1.3 package in R ^106,107^. The second test was a check of the plausibility of the null hypothesis that equal size samples from the transcriptional-adaptation and no-transcriptional-adaptation groups have the same standard deviation. We empirically downsampled (n=49 with percent-positive < 75% in controls, of 55 total genes, same as transcriptional-adaptation group size) the no-transcriptional-adaptation group without replacement, 1,000 times, to generate an empirical null distribution of sample standard deviations from the no-transcriptional-adaptation group. We then compared the observed transcriptional-adaptation group standard deviation to this distribution, and observed that it was among the lowest downsampled values (Figure 5D), suggesting that the transcriptional-adaptation group is unlikely to have a similar sample standard deviation to the no-transcriptional-adaptation group.

### Gene set enrichment analysis

We wondered whether genes involved in any specific biological processes or contexts were overrepresented in the set of genes whose paralogs were significantly upregulated. Therefore, we performed gene set enrichment analysis to check for over-enrichment of any Gene Ontology - Biological Process terms, comparing the following sets of hits against their respective background sets of tested CRISPR targets

1. Bulk RNA-seq CRISPR targets with bootstrap p-value < 0.1, against a background set of all fully analyzed, human genes only.
2. Single-cell RNA-seq CRISPR targets from Frangieh et al., 2021, among those meeting minimum cell count thresholds above, with any paralog in the top-100 largest increase in percent positive list, or if control percent positive > 0.75 with any paralog in top-100 largest increase in mean list, against background of all targets in library
3. Combined (1) and (2) hits against their combined respective backgrounds

We used the clusterProfiler R package v3.12.0 for gene ontology over-representation testing ^48^.

### Networks

Gene regulatory networks are represented as directed graphs. Genes are nodes, and regulatory relationships are edges (e.g., A stimulates B leads to an edge from node A to node B; Figure 3). The biological mechanism presented in recent studies on nonsense-induced transcriptional compensation implies a minimum set of regulatory relationships between an ancestral regulator, its paralog genes, and a downstream target gene ^14,15^. We model gene regulatory networks with, for each gene, two alleles with transcriptional burst activity independent of each other, consistent with observations of transcriptional burst regulation ^108^. The edges between a given regulator gene product and the target gene alleles are set at equal weight, reflecting no regulatory differences at the allele level.

For an upstream regulator gene A that has nonsense-induced transcriptional compensation, there is at least one compensating gene, A’ (referred to as a paralog here). Gene A’ encodes product *A*’. The downstream regulatory target of A and A’ is gene B. Gene B encodes product *B*. Upon mutation of A, the mutant allele of gene A produces product *A*_*nonsense*_ instead of product *A* . During nonsense-induced transcriptional compensation, *A*_*nonsense*_ can regulate alleles of gene A (mutated or not), as well as gene A’, but no longer regulates gene B.

### Core transcriptional bursting model

Our network is built of component genes whose alleles are each modeled with an expanded version of the classic telegraph model, similar to prior work (Figure 3; ^54^). Each allele can reversibly enter an active (“on”; transcribing) or inactive (“off”; quiescent) state, with high or low (by default 0) production rates of that gene’s product, respectively. We assume that any gene product is effectively immediately translated or processed to the relevant functional form capable of regulating a downstream target. In the case of {*A* , *A*’, *B*}, this assumption applies to post-transcriptional regulation, translation, and post-translational processing. In the case of *A*_*nonsense*_, this assumption applies to the hypothesized but unknown mechanisms of nonsense-induced transcriptional compensation. Therefore, for alleles of genes A and A’, and their respective products, there are five consistent reactions:

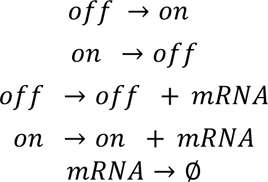

Since for all genes we assume there is no leaky expression in the off state, in the following sections we ignore the possible *off* → *off* + *mRNA* reaction. Alleles of gene B are described by a related but larger set of reactions to reflect the different consequences of regulation by {*A*, *A*’}, and are described below.

As previously described, we make use of reaction propensities under the assumption of the law of mass action, where each propensity function *p*_*i*_(*x*)*dt* gives the probability of reaction *R*_*i*_ occurring in the time step *dt*, for a small *dt*. In the models presented here, gene regulation affects either the reaction rate of an allele entering the active state or entering the inactive state.

### Activation (positive regulation) model

In a model of activating interacting genes in a gene regulatory network, all regulation affects the reaction rate of activation of target alleles. In order to simulate differential effects of A-stimulated B alleles versus A’-stimulated B alleles, we used an expanded gene model for B. The expanded model allows for multiple on-states, corresponding to the different upstream regulators; the different on-states are allowed to have different production rates. These different production rates can reflect biological differences in gene regulation at B loci, including, but not limited to, chromatin changes, differences in transcription factor recruitment, and effects on transcriptional machinery. Therefore, for gene B the full set of reactions is:

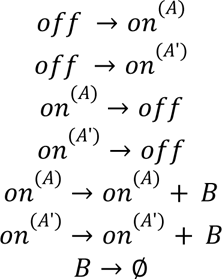

The rates in the reactions described in Figure 3, S4 are:

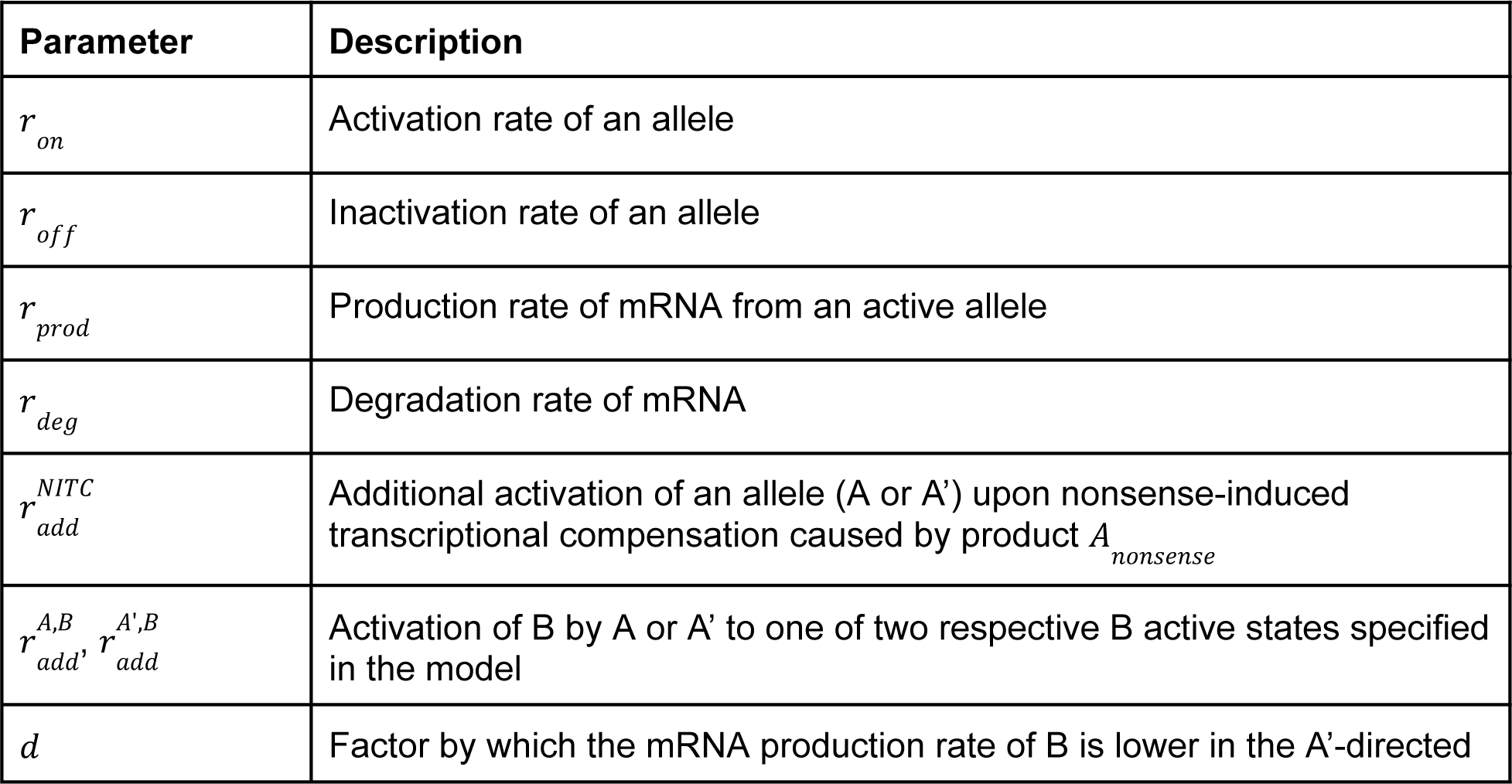
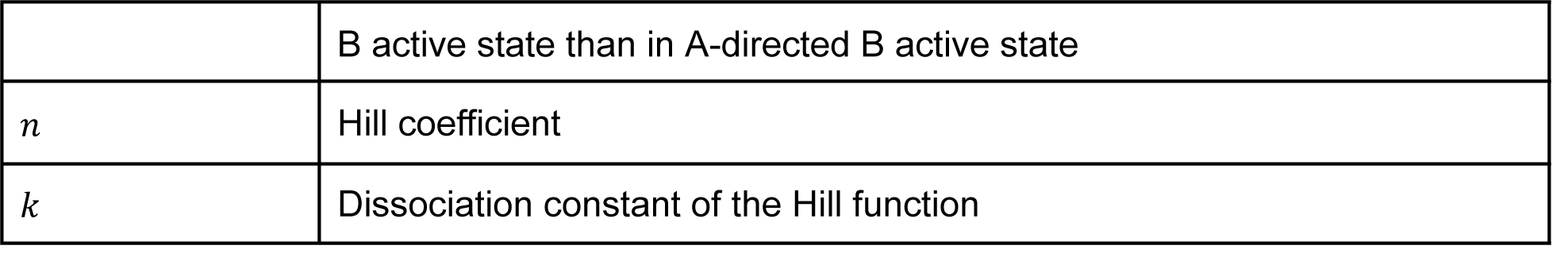

The full model is therefore described as follows. The basal on-rates of A’ and B alleles are assumed to be 0. The mRNA production rate in the off state of all alleles is fixed at 0.

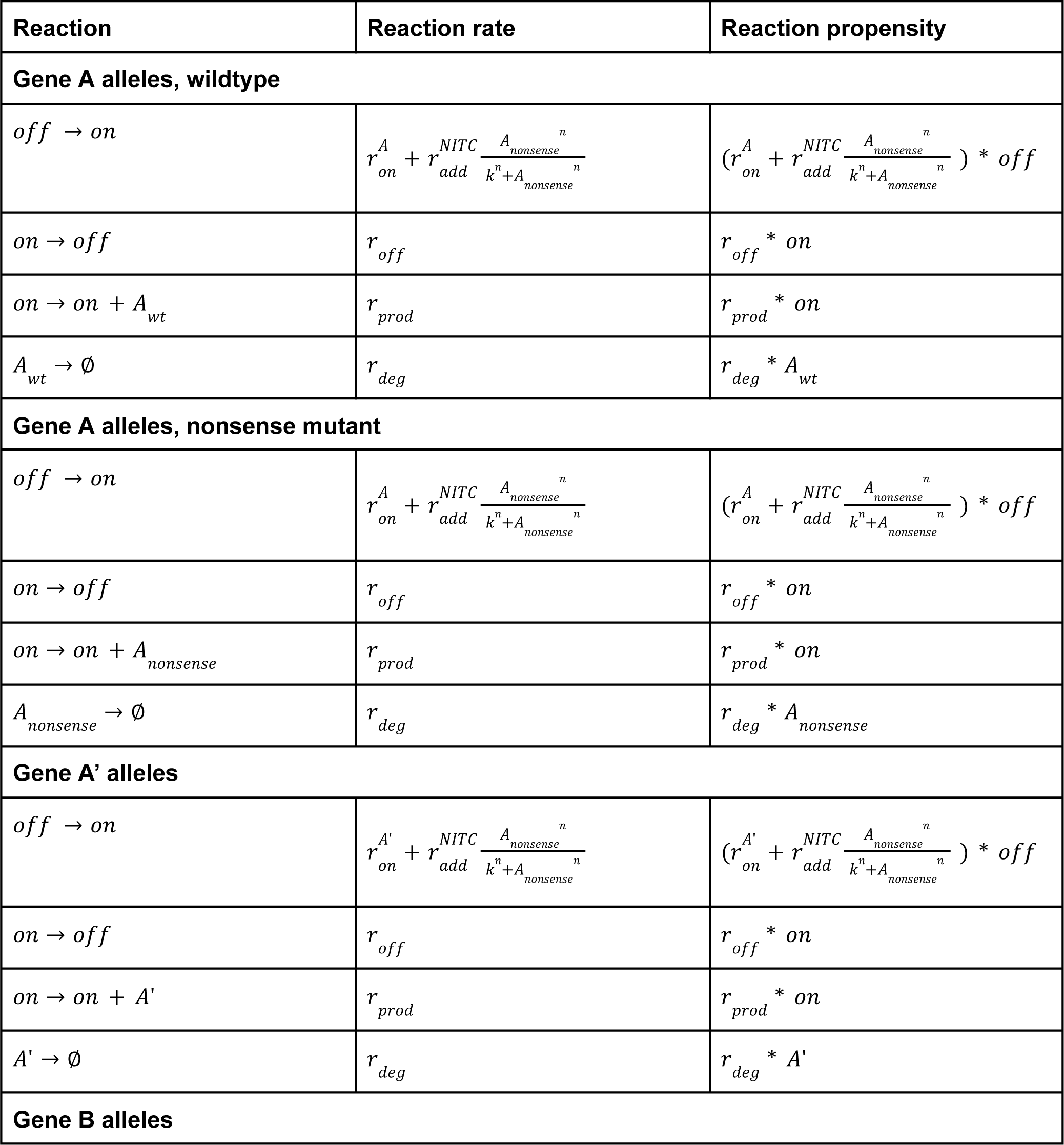
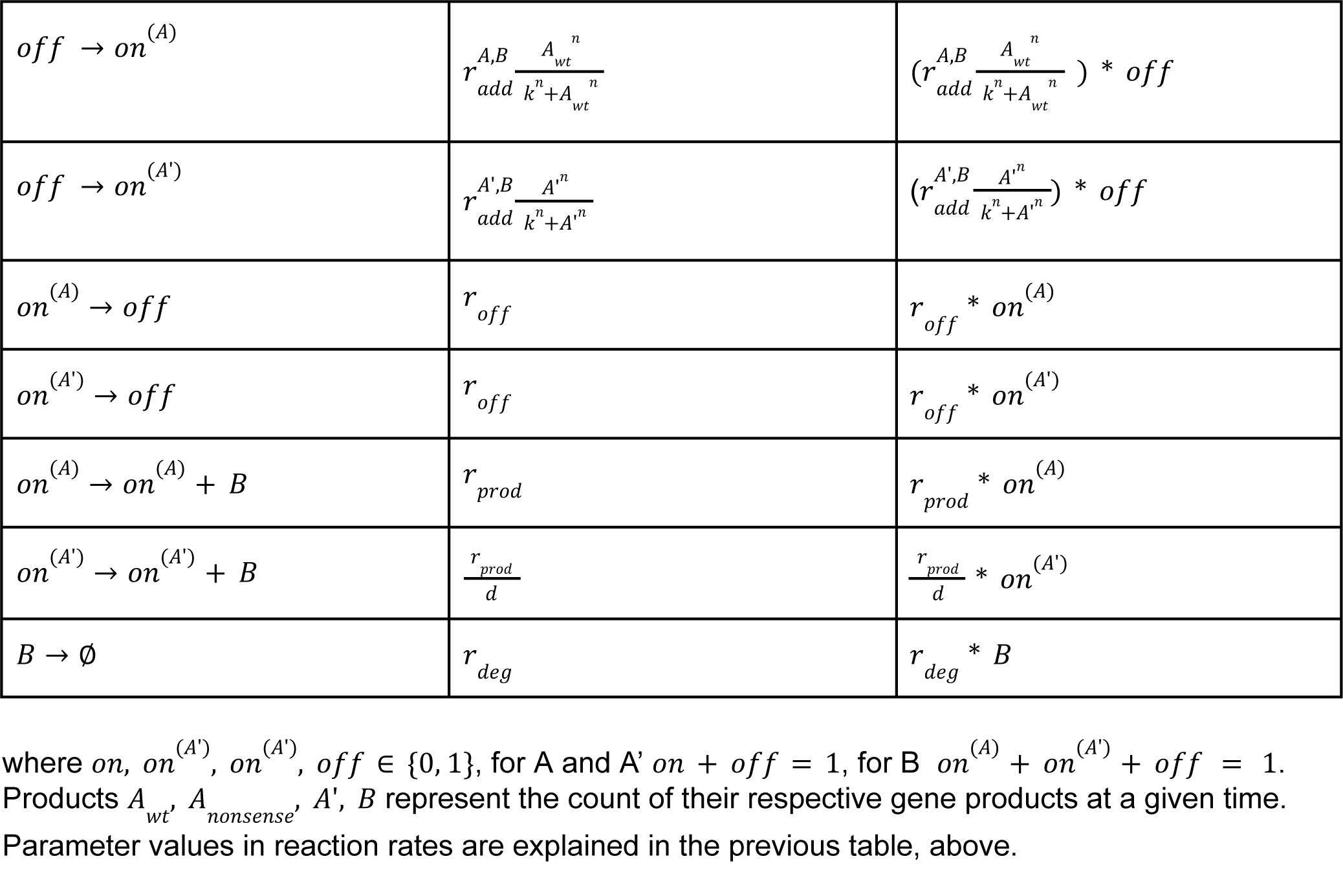

| Reaction | Reaction rate | Reaction propensity |
| --- | --- | --- |
| **Gene A alleles, wildtype** |  |  |
| $off \rightarrow on$ | $r_{on}^A + r_{add}^{NITC} \frac{A_{nonsense}^n}{k^n + A_{nonsense}^n}$ | $(r_{on}^A + r_{add}^{NITC} \frac{A_{nonsense}^n}{k^n + A_{nonsense}^n}) * off$ |
| $on \rightarrow off$ | $r_{off}$ | $r_{off} * on$ |
| $on \rightarrow on + A_{wt}$ | $r_{prod}$ | $r_{prod} * on$ |
| $A_{wt} \rightarrow \emptyset$ | $r_{deg}$ | $r_{deg} * A_{wt}$ |
| **Gene A alleles, nonsense mutant** |  |  |
| $off \rightarrow on$ | $r_{on}^A + r_{add}^{NITC} \frac{A_{nonsense}^n}{k^n + A_{nonsense}^n}$ | $(r_{on}^A + r_{add}^{NITC} \frac{A_{nonsense}^n}{k^n + A_{nonsense}^n}) * off$ |
| $on \rightarrow off$ | $r_{off}$ | $r_{off} * on$ |
| $on \rightarrow on + A_{nonsense}$ | $r_{prod}$ | $r_{prod} * on$ |
| $A_{nonsense} \rightarrow \emptyset$ | $r_{deg}$ | $r_{deg} * A_{nonsense}$ |
| **Gene A' alleles** |  |  |
| $off \rightarrow on$ | $r_{on}^{A'} + r_{add}^{NITC} \frac{A_{nonsense}^n}{k^n + A_{nonsense}^n}$ | $(r_{on}^{A'} + r_{add}^{NITC} \frac{A_{nonsense}^n}{k^n + A_{nonsense}^n}) * off$ |
| $on \rightarrow off$ | $r_{off}$ | $r_{off} * on$ |
| $on \rightarrow on + A'$ | $r_{prod}$ | $r_{prod} * on$ |
| $A' \rightarrow \emptyset$ | $r_{deg}$ | $r_{deg} * A'$ |
| **Gene B alleles** |  |  |
| $off \rightarrow on^{(A)}$ | $r_{add}^{A,B} \frac{A_{wt}^n}{k^n + A_{wt}^n}$ | $(r_{add}^{A,B} \frac{A_{wt}^n}{k^n + A_{wt}^n}) * off$ |
| $off \rightarrow on^{(A')}$ | $r_{add}^{A',B} \frac{A'^n}{k^n + A'^n}$ | $(r_{add}^{A',B} \frac{A'^n}{k^n + A'^n}) * off$ |
| $on^{(A)} \rightarrow off$ | $r_{off}$ | $r_{off} * on^{(A)}$ |
| $on^{(A')} \rightarrow off$ | $r_{off}$ | $r_{off} * on^{(A')}$ |
| $on^{(A)} \rightarrow on^{(A)} + B$ | $r_{prod}$ | $r_{prod} * on^{(A)}$ |
| $on^{(A')} \rightarrow on^{(A')} + B$ | $\frac{r_{prod}}{d}$ | $\frac{r_{prod}}{d} * on^{(A')}$ |
| $B \rightarrow \emptyset$ | $r_{deg}$ | $r_{deg} * B$ |

where $on, on^{(A)}, on^{(A')}, off \in \{0,1\}$ , for A and A' $on + off = 1$ , for B $on^{(A)} + on^{(A')} + off = 1$ . Products $A_{wt}, A_{nonsense}, A', B$ represent the count of their respective gene products at a given time. Parameter values in reaction rates are explained in the previous table, above.

### Multiple-paralog model

Several real genes have multiple annotated paralogs, which in principle could all potentially compensate for mutations in the CRISPR-target gene ^14,15,40^. Therefore, we also developed a gene regulatory network model in which gene A can be compensated for by two paralogs, genes A’1 and A’2 with products *A*’_1_, *A*’_2_ respectively. Similar to the model in the previous section, each allele of A’1 and A’2 are regulated by nonsense-induced transcriptional compensation for A. An expanded model accounting for differences in regulation of B alleles by gene products *A*’_1_, *A*’_2_ leads to 3 possible on states: those directed by *A*_*wt*_, *A*’_1_, *A*’_2_ respectively.

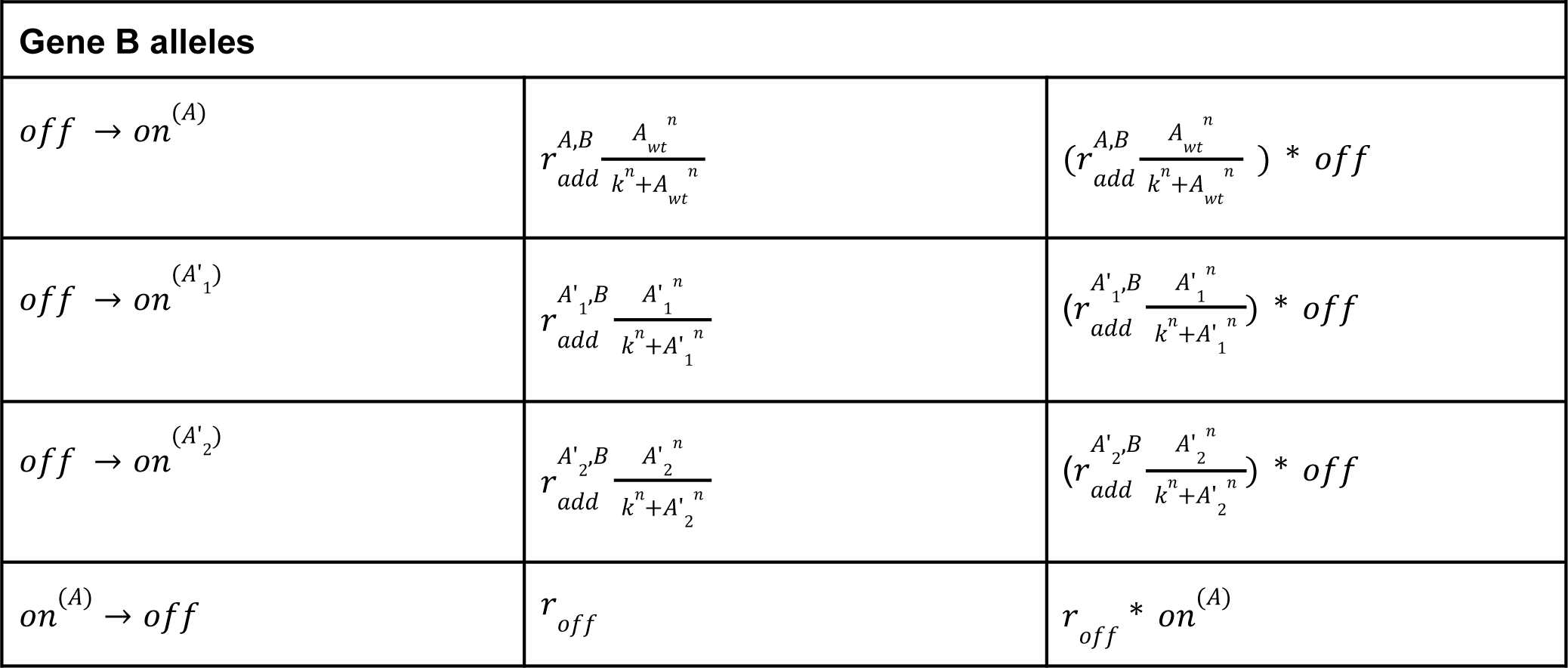
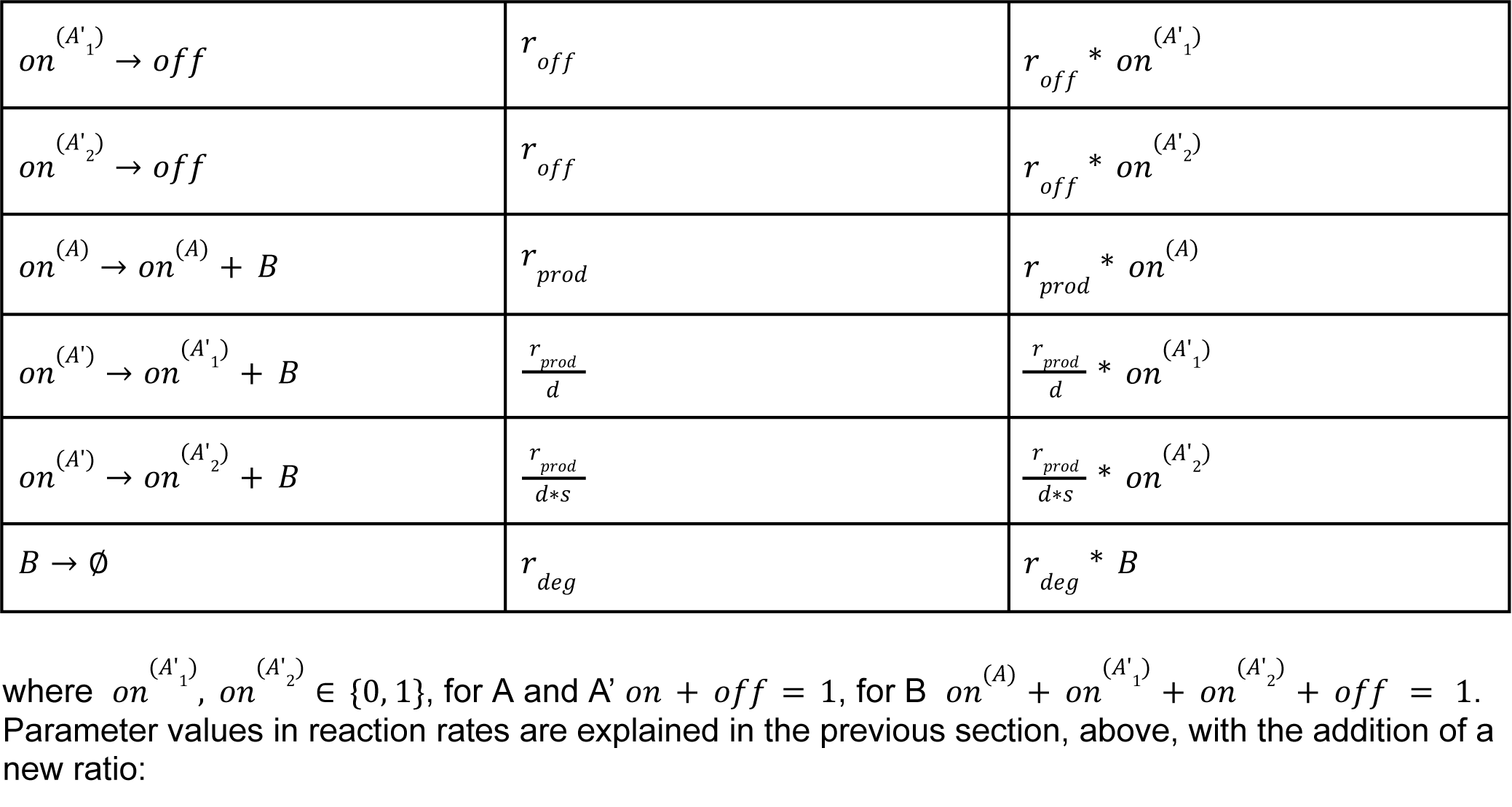

| Gene B alleles |  |  |
| --- | --- | --- |
| $off \rightarrow on^{(A)}$ | $r_{add}^{A,B} \frac{A_{wt}^n}{k^n + A_{wt}^n}$ | $(r_{add}^{A,B} \frac{A_{wt}^n}{k^n + A_{wt}^n}) * off$ |
| $off \rightarrow on^{(A'_1)}$ | $r_{add}^{A'_1,B} \frac{A_1'^n}{k^n + A_1'^n}$ | $(r_{add}^{A'_1,B} \frac{A_1'^n}{k^n + A_1'^n}) * off$ |
| $off \rightarrow on^{(A'_2)}$ | $r_{add}^{A'_2,B} \frac{A_2'^n}{k^n + A_2'^n}$ | $(r_{add}^{A'_2,B} \frac{A_2'^n}{k^n + A_2'^n}) * off$ |
| $on^{(A)} \rightarrow off$ | $r_{off}$ | $r_{off} * on^{(A)}$ |
| $on^{(A'_1)} \rightarrow off$ | $r_{off}$ | $r_{off} * on^{(A'_1)}$ |
| $on^{(A'_2)} \rightarrow off$ | $r_{off}$ | $r_{off} * on^{(A'_2)}$ |
| $on^{(A)} \rightarrow on^{(A)} + B$ | $r_{prod}$ | $r_{prod} * on^{(A)}$ |
| $on^{(A')} \rightarrow on^{(A'_1)} + B$ | $\frac{r_{prod}}{d}$ | $\frac{r_{prod}}{d} * on^{(A'_1)}$ |
| $on^{(A')} \rightarrow on^{(A'_2)} + B$ | $\frac{r_{prod}}{d*s}$ | $\frac{r_{prod}}{d*s} * on^{(A'_2)}$ |
| $B \rightarrow \emptyset$ | $r_{deg}$ | $r_{deg} * B$ |

where $on^{(A'_1)}, on^{(A'_2)} \in \{0, 1\}$ , for A and A' $on + off = 1$ , for B $on^{(A)} + on^{(A'_1)} + on^{(A'_2)} + off = 1$ . Parameter values in reaction rates are explained in the previous section, above, with the addition of a new ratio:

| Parameter | Description |
| --- | --- |
| s | Factor by which the mRNA production rate of B is lower in the A1'-directed B active state than in A'2-directed B active state |

### Repression (negative regulation) model

We also explored gene regulatory network models in which the regulator products *A*, *A*’ are inhibitors of gene B rather than activators. In this model, although nonsense-induced transcriptional compensation continues to activate expression of A and A’, *A*, *A*’ products increase the rate of B allele inactivation instead of activation. To simulate differential effects on B alleles, inspired by differential gene regulatory effects such as chromatin modifications and repressive transcription factor recruitment , we changed the gene model of B to include two off states (one directed by *A* and one by *A*’), with one on state. The two off states could have different rates of reversion to the active state, thereby simulating more or less repressed loci. Therefore, while the model confined to the regulation of genes A and A’ remain the same as above, the model for gene B becomes:

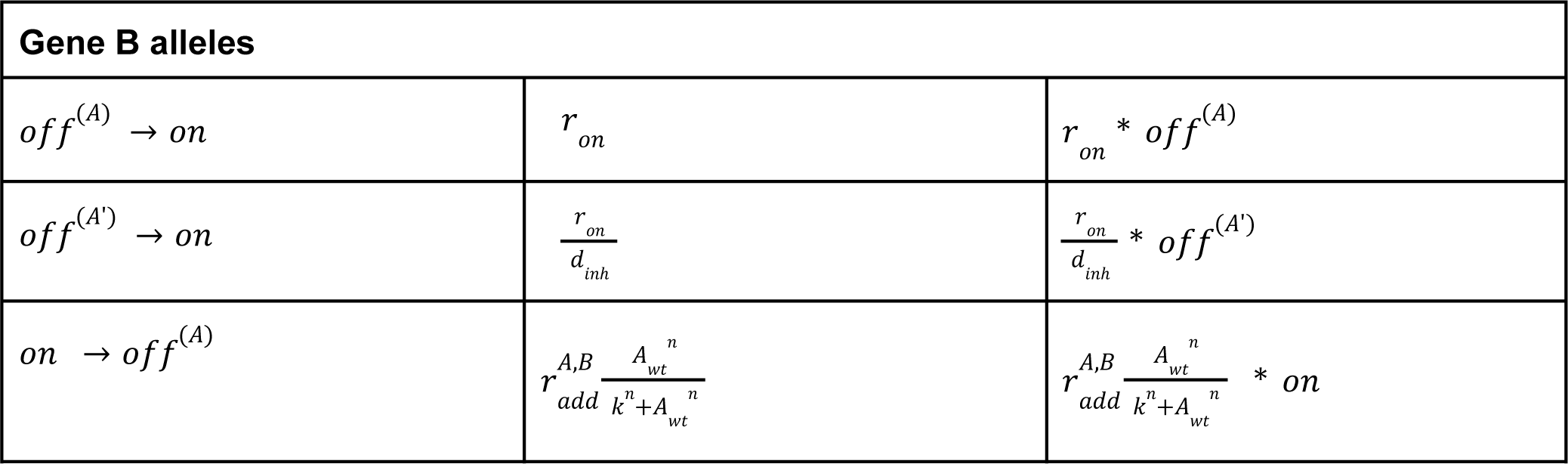
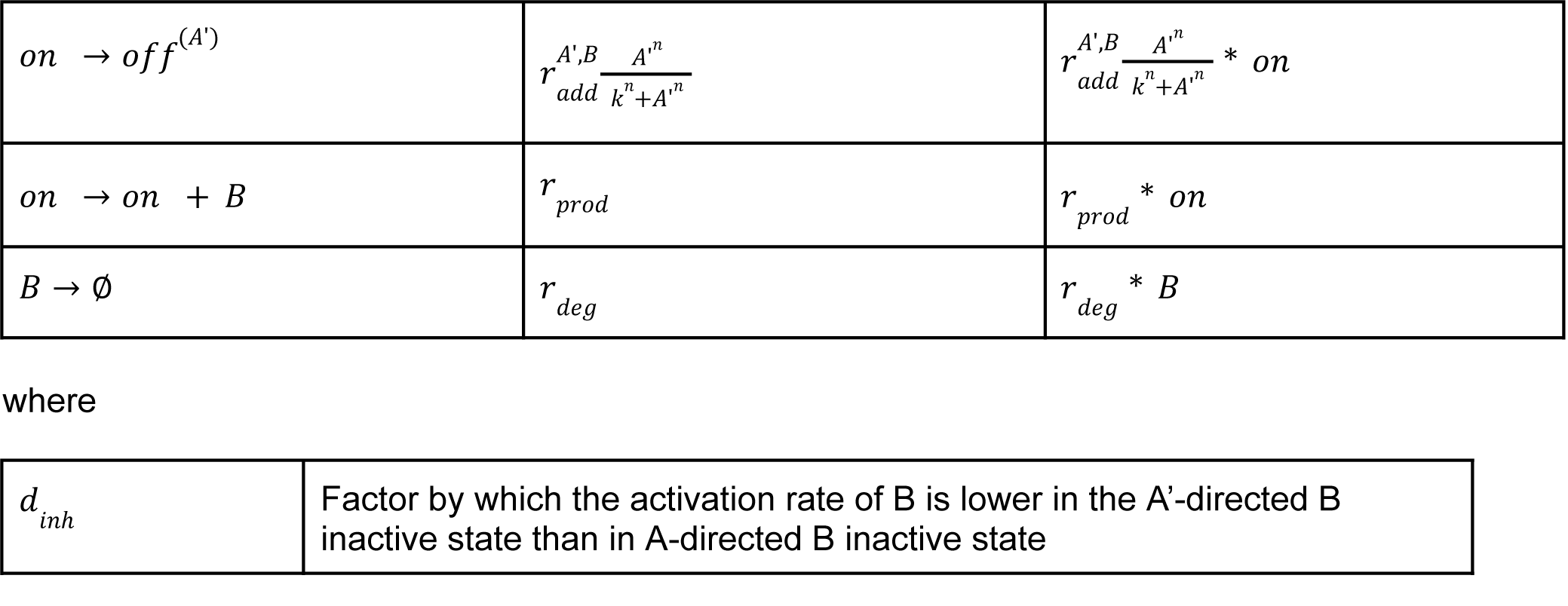

| Gene B alleles |  |  |
| --- | --- | --- |
| $off^{(A)} \rightarrow on$ | $r_{on}$ | $r_{on} * off^{(A)}$ |
| $off^{(A')} \rightarrow on$ | $\frac{r_{on}}{d_{inh}}$ | $\frac{r_{on}}{d_{inh}} * off^{(A')}$ |
| $on \rightarrow off^{(A)}$ | $r_{add}^{A,B} \frac{A_{wt}^n}{k^n + A_{wt}^n}$ | $r_{add}^{A,B} \frac{A_{wt}^n}{k^n + A_{wt}^n} * on$ |
| $on \rightarrow off^{(A')}$ | $r_{add}^{A',B} \frac{A'^n}{k^n + A'^n}$ | $r_{add}^{A',B} \frac{A'^n}{k^n + A'^n} * on$ |
| $on \rightarrow on + B$ | $r_{prod}$ | $r_{prod} * on$ |
| $B \rightarrow \emptyset$ | $r_{deg}$ | $r_{deg} * B$ |

|  |  |
| --- | --- |
| $d_{inh}$ | Factor by which the activation rate of B is lower in the A'-directed B inactive state than in A-directed B inactive state |

### Parameters

We sought to characterize the breadth of possible gene regulatory network outputs given the presence of a transcriptional adaptation regulatory interaction under biologically plausible conditions ^58,60^. Extensive prior research has established transcriptional bursting as a core model of gene expression in eukaryotes. In the bursting model with zero leaky expression as presented here, upon activation of an allele the steady state expected mRNA abundance derived from that allele rises from 0 to *r*_*prod*_ /*r*_*deg*_. When mRNA abundance is high enough to exceed a threshold specified by the dissociation constant of the Hill function (*k*) above, there is a higher probability of activation of alleles of downstream targets, further modulated by *r_add_* parameters, above. The dissociation constant is defined as:

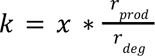

where x is the fraction of the active-steady-state expression level required to be exceeded for regulation of the target allele. In our simulations, we use *x* = 0. 5 to represent regulation that can occur with some degree of expression of an upstream regulator, so that regulation is not presumed to require prolonged active high-expression states, consistent with published burst duration measurements ^58^. Other studies have also simulated transcriptional bursts in gene regulatory networks based on these parameter ranges ^60^.

Published telegraph model inferred parameter ranges based on allele-resolved single-cell RNA sequencing data provide a useful guide to ranges for the core allele-level parameters ^58^. We reanalyzed the data in Larsson et al., 2019, to summarize, for each reported gene expressed in mouse fibroblasts, relative to the degradation rate (*r*_*deg*_ arbitrarily fixed at 1), what are the inferred values of *r*_*on*_, *r*_*off*_, *r*_*prod*_. In order to preserve overall burst frequency and duration ranges, we focused on the calculated values of: 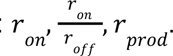 Based on the observed results, we conducted simulations over parameter ranges spanning approximately 2-3 orders of magnitude:

| Parameter | Meaning | Lower limit | Upper limit |
| --- | --- | --- | --- |
| $r_{deg}$ | degradation rate of mRNA | 1 | 1 |
| $r_{prod}$ | production rate of mRNA in on state | 1 | 1000 |
| $r_{on}$ | activation rate of an allele (by default, an A allele) | 0.1 | 10 |
| on/off ratio | $\frac{r_{on}}{r_{off}}$; the factor by which the rate of inactivation is lower than activation | 0.01 | 2 |

Consistent with prior studies, we also explored a range of values of Hill coefficient *n*, to ensure that we adequately sampled over different steepness levels of the Hill function representing regulatory relationships ^54^.

| Parameter | Meaning | Lower limit | Upper limit |
| --- | --- | --- | --- |
| Hill coefficient | $n$; the Hill coefficient. How “switch-like” regulatory effects are. | 0.1 | 5 |

Extending the basic allele-level telegraph model to our nonsense-induced transcriptional compensation gene regulatory network model, we needed to pick reasonable ranges for other parameters. For the primary simulations discussed in Figures 3-4, we assumed zero basal activation of paralog A’ alleles. We focused on characterizing the relative strengths of interactions, as that directly reflects the differences between a regulator, its paralog, and their downstream target rather than absolute simulation parameter values. Therefore, in order to explore network outputs over similarly large ranges of other interaction strengths (roughly two orders of magnitude), and to ensure that activation rates were at least partially overlapping with the observed rates from Larsson et al., 2019, we further considered rate ratios as follows:

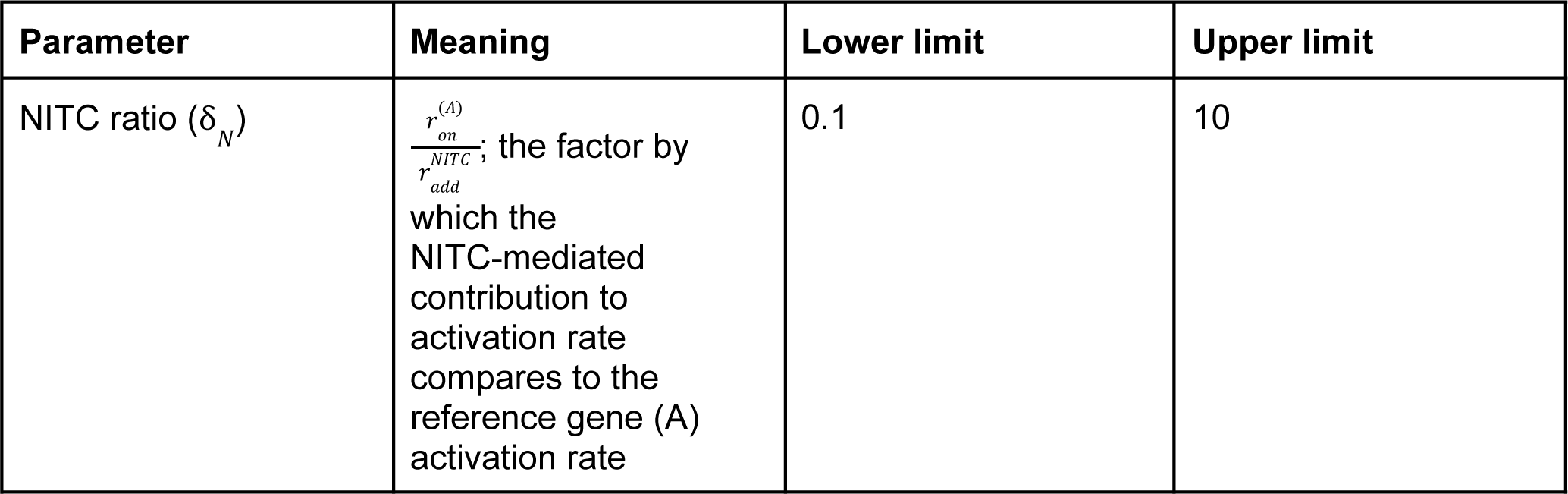

For each of the different models, we sampled over several other rate and ratio model parameters.

In the base case, a model with one paralog and stimulating regulation of gene B, we also considered:

For each of the different models, we sampled over several other rate and ratio model parameters. In the base case, a model with one paralog and stimulating regulation of gene B, we also considered:
| Parameter | Meaning | Lower limit | Upper limit |
| --- | --- | --- | --- |
| $r_{add}^{A,B}$ | Contribution to activation of B by A to A-directed B active state | 0.1 | 10 |
| A/A' activation ratio ($\delta_A$) | $\frac{r_{add}^{A,B}}{r_{add}^{A',B}}$; the factor by which A is more effective at activating B alleles to A-directed active state than A' to A'-directed active state | 0.1 | 10 |
| A/A' production ratio ($\delta_p$) | $d$; the factor by which the mRNA production rate of B is lower in the A'-directed B active state than in A-directed B active state | 0.1 | 10 |

In a set of simulations of the stimulatory model including non-zero paralog A’ expression at baseline, we also sampled over:

| Parameter | Meaning | Lower limit | Upper limit |
| --- | --- | --- | --- |
| Basal A' ratio | $\frac{r_{on}^{(A)}}{r_{on}^{(A')}}$; the factor by which A allele activation rates are higher than A' alleles at baseline | 1 | 100 |

In the model expanded to consider multiple paralogs, we conducted simulations with and without basal paralog expression. In one set of simulations, we fixed the effects of both paralogs, A’1 and A’2, on B, to be equal. In another set of simulations, we sampled over values of a new ratio describing how expression varies between A’1-directed and A’2-directed B-active states.

| Parameter | Meaning | Lower limit | Upper limit |
| --- | --- | --- | --- |
| A'1/A'2 prod ratio ($\delta_R$) | $s$; Factor by which the mRNA production rate of B is lower in the A'1-directed B active state than in A'2-directed B active state | 1 | 100 |

In a model of inhibitory regulation of gene B, instead of A/A’ add-on ratio and A/A’ prod ratio, we sampled over parameter ratios: A/A’ add-off ratio and B,A/B,A’ on ratio, to reflect the regulator contributions to inactivation rates to their respective B off states and their respective off states’ rates of activation (to a common single active state; see Figure S14), which could differ.

| Parameter | Meaning | Lower limit | Upper limit |
| --- | --- | --- | --- |
| $r_{add,off}^{A,B}$ | Contribution to inactivation of B by A to A-directed B inactive state | 0.1 | 10 |
| A/A' add-off ratio | $\frac{r_{add,off}^{A,B}}{r_{add,off}^{A',B}}$; the factor by which A is more effective at inactivating B alleles to A-directed inactive state than A' to A'-directed inactive state | 0.1 | 10 |
| A/A' B-activation ratio | $d_{inh}$; factor by which the activation rate of B is lower in the A'-directed B inactive state than in A-directed B inactive state | 0.1 | 10 |

The parameter sets (spanning 8 or 9 parameters or parameter ratios, depending on the model and assumptions) for each set of simulations are described in Table S4.

For each primary set of simulations, we used latin hypercube sampling to homogeneously sample over the multidimensional parameter spaces. In each case we drew 10,000 parameter sets from within the upper and lower boundaries of each log_10_-transformed parameter range (https://www.mathworks.com/matlabcentral/fileexchange/45793-latin-hypercube). Log transformation was used to more evenly sample over orders of magnitude.

### Simulations

We simulated each of the 3 network models under at least 2 different assumed conditions related to differential effects or paralog expression (see Figures), for approximately 10,000 parameter sets, resulting in a total of approximately 60,000 simulations across 3 network models, each containing consecutive time periods simulating 3 genotypes: wildtype (neither A allele mutated), heterozygous (one A allele mutated), and homozygous-mutant (two A alleles mutated). We also conducted parameter subspace resampling for smaller numbers of parameter sets. We used Gillespie’s next reaction method, as previously described ^32,54,57^. We computed for a total of 300,000 timesteps per simulation, i.e., 100,000 in each genotype. In each simulation, at time t=0 all alleles were in the inactive state and the mRNA count was fixed at 1 for *A* and 0 for all other products. We implemented the simulations in MATLAB R2017a and R2021b ^54^. Each simulation took between 30 seconds and 12 minutes to run, depending on the parameter values, leading to a total simulation time of approximately two weeks using 8 cores running in parallel.

### Pseudo-single-cell analysis and autocorrelation

In order to simulate snapshot single-cell population measurements of gene expression from these simulations, we split the simulation traces into 300-timestep-unit segments and used the first set of values (of DNA activation states and product abundances) as a sampled “pseudo-single-cell” measurement, as discussed in prior work ^54^. We needed to confirm that our sub-simulation samples were not susceptible to unexpected autocorrelation that might interfere with using these samples as independent pseudo-random samples of single cells drawn from the underlying distributions emergent from the network simulations. Therefore, we used the stats::acf function in R to show that 100-step lags were within the 95% confidence intervals for random autocorrelation for a random sample of parameter sets, and set the lag at an even higher value, 300, out of an abundance of caution given the range of possible parameter combinations.

### Steady state analysis

We also wanted to confirm that the output of our numerically simulated stochastic network models fitted with other ways of estimating the outputs of these same networks, both as a quality control and to highlight the added benefits related to simulating variability with stochastic simulations in studies of complex networks. Therefore, we used the ode45 solver in MATLAB R2017a to deterministically estimate the steady state outputs of all gene product levels in each genotype for 100 parameter sets. The systems of differential equations for each model included 18, 25, and 18 equations each, for the single-paralog stimulation, two-paralog stimulation, and single-paralog repression models, respectively. Initial conditions included all alleles set to the off state and all mRNA levels set to 0. A timespan of 500 units was used, and a random sample of results were inspected to ensure that mRNA level estimates had reached steady state. We then compared these estimated steady state outputs to the pseudo-single-cell population means from our simulations, and observed very high concordance at the absolute abundance level. The simulations in which a gene product did not have a pseudo-single-cell mean similar to the ode45 steady state solution were most often for parameter sets with high Hill coefficients (*n*), reflecting high non-linearity in regulatory interactions.

### Distribution shape statistics

We sought to describe the variability in gene expression emerging from gene regulatory networks with transcriptional adaptation, and to quantify differences in aspects of variability between network outputs given different parameter values. Therefore, we calculated several summary statistics related to distribution shape to highlight important features of gene expression distributions.

For each gene product in each genotype, we calculated the first four empirical moments of gene expression distributions which describe different aspects of distribution shape. Briefly, the first moment is mean (µ), which would parallel steady state output for a symmetric unimodal distribution. The second moment is variance, for an overall estimate of spread in the distribution. Instead of variance (σ^2^) specifically, we focus our analyses instead on the coefficient of variation 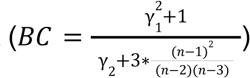, which moredirectly parallels the percentage of spread relative to the mean. The third moment is skewness (γ ),which will be positive for right-skewed distributions and negative for left-skewed distributions. The fourth moment is kurtosis, specifically here the excess kurtosis, (γ ), which, among other uses, is positively correlated with the heaviness of a distribution’s tails.

We also calculated several additional statistics related to distribution shape. Particularly for follow-up analyses, we computed the bimodality coefficient 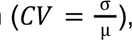 ^109,110^. The bimodality coefficient has a number of useful features. It is a statistic with value constrained to [0, 1]. Uniform distributions will have 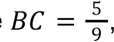, while unimodal distributions will have values closer to 0 and bimodal (or multimodal) distributions will have values closer to 1. Two additional statistics associated with distribution shape are Gini coefficient and entropy. Gini coefficient is constrained to [0, 1], where 1 corresponds to a distribution in which one cell has non-zero expression and all others have zero expression, and 0 corresponds to a distribution in which all cells express the same amount. We calculated entropy over the binned expression axis, considering 30 bins spread evenly across the range of expression values. As described in Results, we used these distribution shape statistics both directly in analyses of model parameter effects as well as indirectly in a shape classifier algorithm, below, which we then also used for analyses of model parameter effects on gene expression.

### Normalized distribution shape statistics

Initial exploration of the distribution statistics revealed that several systematically correlated, often nonlinearly, with the overall sample mean. Therefore, to partially correct for differences in mean explaining differences in other statistics, we performed local regression (LOESS) of each statistic against mean, using default settings in the loess function in R with span = 0.1. The residual of the observed minus LOESS fitted statistic value at an observation’s same mean can be considered as a mean-corrected version of the observed statistic. For *BC*, the LOESS residual is called *BC*^*res*^. One such LOESS-corrected statistic (bimodality coefficient) was used in the distribution shape classifier below, in conjunction with the uncorrected statistic value.

### Distribution shape classification

We present several analyses centered on the question of when an expression distribution can remain robust to the mutation of an upstream regulator. Therefore, we built an algorithm for classifying distribution shapes to reflect plausibly important differences. We were particularly interested in a robust method for identifying whether a distribution was unimodal and symmetric, suggesting a degree of homogeneity in expression. For distributions that were bimodal (or multimodal), one could imagine different emergent properties in a population of cells, e.g., with bistability or other kinds of functional diversity. For distributions that were unimodal but not symmetric, i.e., skewed, one could imagine a bias toward low-frequency diversity in behavior, either being very high expressors or very low expressors. Lastly, we also needed to identify when expression levels were very low in general, reflecting overall minimal transcriptional activity.

The algorithm sorts distributions into 1 of 5 classes:

1. Low-expression
2. Unimodal symmetric
3. Right-skewed unimodal
4. Left-skewed unimodal
5. Bimodal (or multimodal)

It starts by considering whether µ < 10. If so, the distribution is called low-expression. Next, if *BC* > 0. 555 and if *BC*^*res*^ > 0. 1, the distribution is called bimodal (or multimodal). After that, if γ > 1, then the distribution is called right-skewed unimodal and if γ < (− 1), then the distribution is called left-skewed unimodal. All remaining distributions have relatively high expression, low absolute skewness, and low bimodality coefficient, and they are called unimodal symmetric. This algorithm was the result of numerous modified iterations during algorithm development, paired with manual inspections of classifier results on random selections of hundreds of simulation results. Classifier result accuracy (i.e., whether a distribution classification by the algorithm is in agreement with manual assignment) was high (greater than 80%, often greater than 90%) for all distribution shape classes.

We focused our analyses on whether or not a distribution was called unimodal symmetric and whether the unimodal symmetric class could be made robust to mutation of an upstream regulator.

### Decision tree analysis

We wanted to check whether there were subspaces of parameter space that are enriched for gene regulatory outputs that display behavior of interest, e.g., robustness of shape to mutation of an upstream regulator, for unimodal symmetric distributions. Therefore, as previously described, we trained decision tree classifiers on simulation results paired with algorithm-assigned distribution shape classes, particularly for gene product *B*. According to the binary classifications described in the respective sections of Results, we performed decision tree optimization in R using partykit v1.2-16 and its associated dependencies, with alpha = 0.01 for variable selection. As discussed in the respective sections of Results, we validated decision tree results by resampling parameter sets from the parameter subspaces bounded by the decision rules enriching for the specific behavior of interest. In each case we used latin hypercube sampling, as described above, to sample 100 parameter sets from each subspace and conducted simulations, also as described above.

### Statistical analysis

Unless noted otherwise in figure legends, error bars represent standard error of the mean. RNA-seq data analysis, including bootstrap resampling, was performed in Python v3.9.15 using gprofiler-official v1.0.0, matplotlib v3.6.2, numpy v1.23.5, pandas v1.5.2, pydeseq2 v0.3.5, requests v2.28.1, scipy v1.9.3, and seaborn v0.12.1. Latin hypercube sampling and simulations were run in MATLAB R2017a and R2021b. All remaining statistical analysis and graph generation was performed in R v3.6.1 using packages readxl v1.4.0, partykit v1.2-16, mvtnorm v1.1-3, libcoin v1.0-9, ggalluvial v0.12.3, entropy v1.3.1, svglite v2.1.0, corrplot v0.92, ggrepel v0.9.1, e1071 v1.7-11, diptest v0.76-0, gridExtra v2.3, Hmisc v4.7-0, Formula v1.2-4, survival v3.3-1, lattice v0.20-45, ineq v0.2-13, magrittr v2.0.3, forcats v0.5.1, stringr v1.4.0, dplyr v1.0.9, purrr v0.3.4, readr v2.1.2, tidyr v1.2.0, tibble v3.1.7, ggplot2 v3.3.6, tidyverse v1.3.1.

### Data and code availability

GEO accession numbers for publicly accessible, previously published RNA-seq datasets analyzed here are noted in Table S1. Processed Perturb-seq data tables are available at https://singlecell.broadinstitute.org/, accession SCP1064, “Control” condition. Simulation outputs analyzed in this paper are available for download on Google Drive (https://drive.google.com/drive/folders/1JN8P7zWe03uO3oDxoz4RZadKfz8Ar2kX?usp=sharing). All code required to reproduce the figures in this paper is available on github (https://github.com/GoyalLab/TA_prevalence_constraints_public)

## Supplementary Figure Legends

S1: Supplementary analysis of paralog upregulation after CRISPR/Cas9-mediated knockouts.

A. Per-knockout-target paralog differential expression results using alternative differential expression inclusion criteria to Figure 1C. Differential expression counted if log2 fold-change > 0.5, adjusted p-value < 0.05, and if basemean > 10. Per knockout target: expected frequency of paralog upregulation on the x axis, observed frequency of paralog upregulation on the y axis. Error bars represent the standard error of the null expected mean frequency of paralog upregulation across 10000 bootstrap samples (see Methods).

B. Per-knockout-target paralog differential expression results using alternative differential expression inclusion criteria to Figure 1C. Differential expression counted if log2 fold-change > 0 and if adjusted p-value < 0.05. Per knockout target: expected frequency of paralog upregulation on the x axis, observed frequency of paralog upregulation on the y axis. Error bars represent the standard error of the null expected mean frequency of paralog upregulation across 10000 bootstrap samples (see Methods).

C. Reanalysis of RNA-seq from ^14^ after three knockouts (see reference manuscript’s Figure 4A) using the paralog upregulation pipeline presented here. Error bars represent standard deviation of bootstrap samples, in gray. Observed fractions of paralogs upregulated, in purple.

D. Perturb-seq-based single-cell paralog expression change after reference gene knockout. Per paralog, percentage of cells positive for that paralog expression in non-template control guide-treated cells on x-axis, percentage of cells positive for that paralog expression in cells treated with guides targeting a reference CRISPR target for which the gene is a paralog.

Paralogs ranked in the top-100 of absolute increases per quantification method marked in magenta. All paralogs of all knockouts shown, using cells meeting UMI count inclusion criteria, regardless of whether minimum cell count per guide inclusion criteria were met, listed in Methods.

S2: Analysis of transcriptional adaptation-associated genes’ expression levels with knockout target paralog upregulation

A. Expression levels (DESeq2 basemean) of COMPASS complex and Upf genes important for nonsense-mediated decay in humans, for CRISPR targets in GSE151825 displaying paralog upregulation by transcriptional adaptation (“TA”) or not displaying paralog upregulation by transcriptional adaptation (“non-TA”).

B. Expression levels (DESeq2 basemean) of COMPASS complex and Upf genes important for nonsense-mediated decay in mice, for CRISPR targets in GSE145653-1 displaying paralog upregulation (with adaptation) or not displaying paralog upregulation (without adaptation).

S3: Analysis of whether there are paralog and target gene feature associations with expression change after reference gene knockout

A. Per-knockout-target scatter plots of each paralog’s sequence homology with the knockout target versus log_2_ fold-change of paralog after reference gene knockout.

B. Per-knockout-target scatter plots of each knockout target gene length with the paralog upregulation frequency after knockout.

S4: Simulation transcriptional burst kinetics summary and resulting parameter set classifications, before and after mutation

A. Parameter descriptions for the model described in Figure 3A.

B. From ^58^ Supplementary Table 1, for mouse fibroblasts transcriptome-wide, histograms of (left), burst on-rates,

C. ratios, per allele, of burst on to burst off rates, and

D. mRNA production rates

E. Prevalence of distribution classes for gene product B across the sampled parameter space for 10000 sampled parameter sets. Sankey plot demonstrates classes of gene B in the wildtype A genotype and, after mutation, in the heterozygous A genotype. Height of each section proportional to number of classified parameter sets.

S5: Autocorrelation analysis of simulated gene expression traces

A. Three random, representative autocorrelation plots for per-100-timestep samples from gene expression traces in the wildtype genotype. Dashed red line represents the 95% confidence bound around zero autocorrelation.

B. Equivalent autocorrelation plots in the heterozygous mutant genotype.

C. Equivalent autocorrelation plots in the homozygous mutant genotype.

S6: Steady-state solutions of gene regulatory network systems of differential equations

A. For each gene product, steady-state solutions of the full system of ordinary differential equations representing the network in Figure 2A, see Methods, in the wildtype genotype, compared against simulation average expression levels.

B. Equivalent plots in the heterozygous genotype.

C. Equivalent plots in the homozygous mutant genotype.

S7: Summary statistic covariation with mean

A. Bimodality coefficient against log(mean) for gene product B in the heterozygous genotype, Red dashed line marks mean = 10.

B. Entropy against log(mean) for gene product B in the heterozygous genotype, Red dashed line marks mean = 10.

C. Coefficient of variation against log(mean) for gene product B in the heterozygous genotype, Red dashed line marks mean = 10.

D. Skewness against log(mean) for gene product B in the heterozygous genotype, Red dashed line marks mean = 10.

E. Gini coefficient against log(mean) for gene product B in the heterozygous genotype, Red dashed line marks mean = 10.

S8: Distribution shape classifier and model parameter associations with classifier component statistics

A. Random samples of distributions assigned to each class were sampled (20-40 per class) and manually assessed for whether they matched class assignment. Fraction matching each class assignment on y-axis.

B. Scatter plots and correlation between selected summary statistics of B distributions in the heterozygous genotype and model parameters. Red arrows highlighting density of observations, see correlation reported in Results.

S9: Distribution class results with basal paralog expression

A. Prevalence of distribution classes for gene product B across the sampled parameter space for 10000 sampled parameter sets, including some (variable) basal paralog expression. Sankey plot demonstrates classes of gene B in the wildtype A genotype and, after mutation, in the heterozygous A genotype.

S10: Decision tree analysis of distribution shape in the heterozygous genotype

A. Decision tree trained on all parameter sets, on whether gene B is classified as unimodal symmetric shape in the heterozygous genotype.

S11: A gene regulatory network with transcriptional adaptation by multiple paralogs

A. Schematic of a gene regulatory network with transcriptional adaptation to mutation, with two paralogs. Two alleles of each gene, with bursty transcription of gene products at each allele. A mutated reference gene, A, regulates downstream effector gene B. When mutated, nonsense copies of A product stochastically upregulate paralogs of A, called A’1 and A’2. A’1 and A’2 can also regulate B, albeit with different strengths than A. In one set of simulations, A’1 and A’2 have the same parameter values. In a second set of simulations, they have different effects on the production rate of B. Hill functions are used in propensities for regulatory relationships between gene products and target alleles. See Methods for full model specification.

B. Prevalence of distribution classes for gene product B across the sampled parameter space for 10000 sampled parameter sets. Both paralogs have the same parameter values. Sankey plot demonstrates classes of gene B in the wildtype A genotype and, after mutation, in the heterozygous A genotype.

C. Prevalence of distribution classes for gene product B across the sampled parameter space for 10000 sampled parameter sets. A’1 and A’2 have different effects on the production rate of B. Sankey plot demonstrates classes of gene B in the wildtype A genotype and, after mutation, in the heterozygous A genotype.

S12: Decision tree analysis for robustness of unimodal symmetric B expression in a multi-paralog model with equal paralog parameters

A. Decision tree trained on parameter sets in which gene B is classified as unimodal symmetric in the wildtype phenotype, on whether gene B is classified as having unimodal symmetric shape in the heterozygous genotype.

S13: Decision tree analysis for robustness of unimodal symmetric B expression in a multi-paralog model with different paralog parameters

A. Decision tree trained on parameter sets in which gene B is classified as unimodal symmetric in the wildtype phenotype, on whether gene B is classified as having unimodal symmetric shape in the heterozygous genotype.

S14: A gene regulatory network with transcriptional adaptation for a repressor

A. Schematic of a gene regulatory network with transcriptional adaptation to mutation, in which the regulatory interactions between A and B are repressive.Two alleles of each gene, with bursty transcription of gene products at each allele. A mutated reference gene, A, regulates downstream effector gene B. When mutated, nonsense copies of A product stochastically upregulate a paralog of A, called A’. A’ can also regulate B, with different parameters than A. In one set of simulations, A’ does not have any basal expression (without the effects of NITC); in another set of simulations, A’ has variable amounts of basal expression. Hill functions are used in propensities for regulatory relationships between gene products and target alleles. See Methods for full model specification.

B. Prevalence of distribution classes for all gene products across the sampled parameter space for 10000 sampled parameter sets. No basal expression of A’. Sankey plot demonstrates classes of each gene in the wildtype A genotype and, after mutation, in the heterozygous A genotype, and after a second mutation, in the homozygous mutant A genotype.

C. Prevalence of distribution classes for each gene product across the sampled parameter space for 10000 sampled parameter sets. Variable amounts of basal expression of A’. Sankey plot demonstrates classes of gene B in the wildtype A genotype and, after mutation, in the heterozygous A genotype.

S15: Decision tree analysis for robustness of unimodal symmetric B expression in a network model of a repressor, without basal paralog expression

A. Decision tree trained on parameter sets in which gene B is classified as unimodal symmetric in the wildtype phenotype, on whether gene B is classified as having unimodal symmetric shape in the heterozygous genotype.

S16: Decision tree analysis for robustness of unimodal symmetric B expression in a network model of a repressor, with basal paralog expression

A. Decision tree trained on parameter sets in which gene B is classified as unimodal symmetric in the wildtype phenotype, on whether gene B is classified as having unimodal symmetric shape in the heterozygous genotype.

## Supplementary Tables

S1: Bulk RNA-seq datasets reanalyzed in this study. GEO entries, experimental designs, and descriptions of knockout targets are provided.

S2: Bulk RNA-seq-based knockout targets analyzed for paralog upregulation.

S3: Perturb-seq-based knockout target-paralog pairs with signs of potential paralog upregulation. Target-paralog pairs included if the paralog was among either in the top-100 increases in percent-positive cells (for paralogs below 75% positive in controls) or in the top-100 increases in mean expression level (for paralogs above 75% positive in controls).

S4: Latin hypercube sampled parameter sets for simulations presented in this study.

