## Supplementary Figures for "Prevalence of and gene regulatory constraints on transcriptional adaptation in single cells"

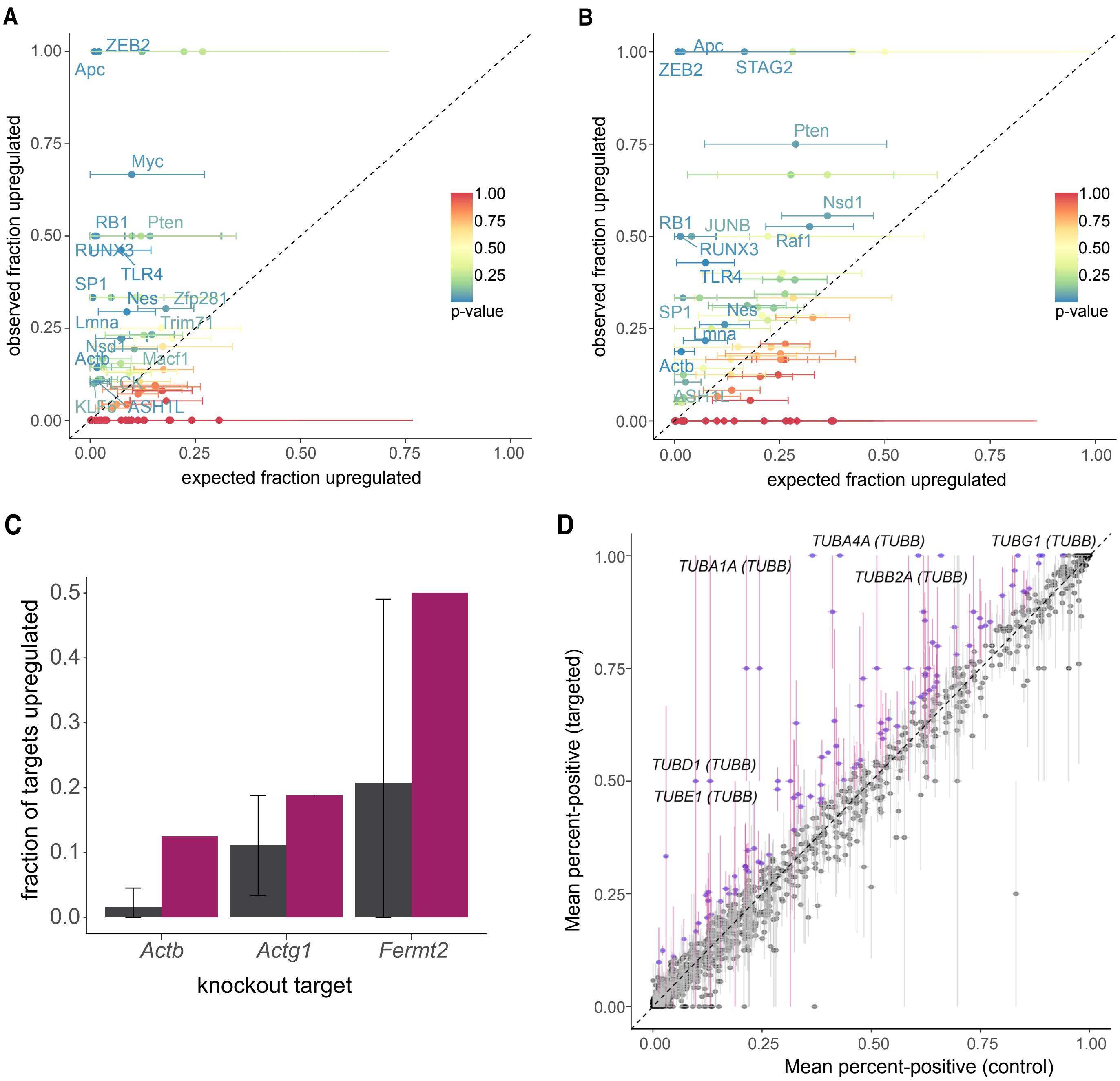


**Supplementary Figure 1**

A

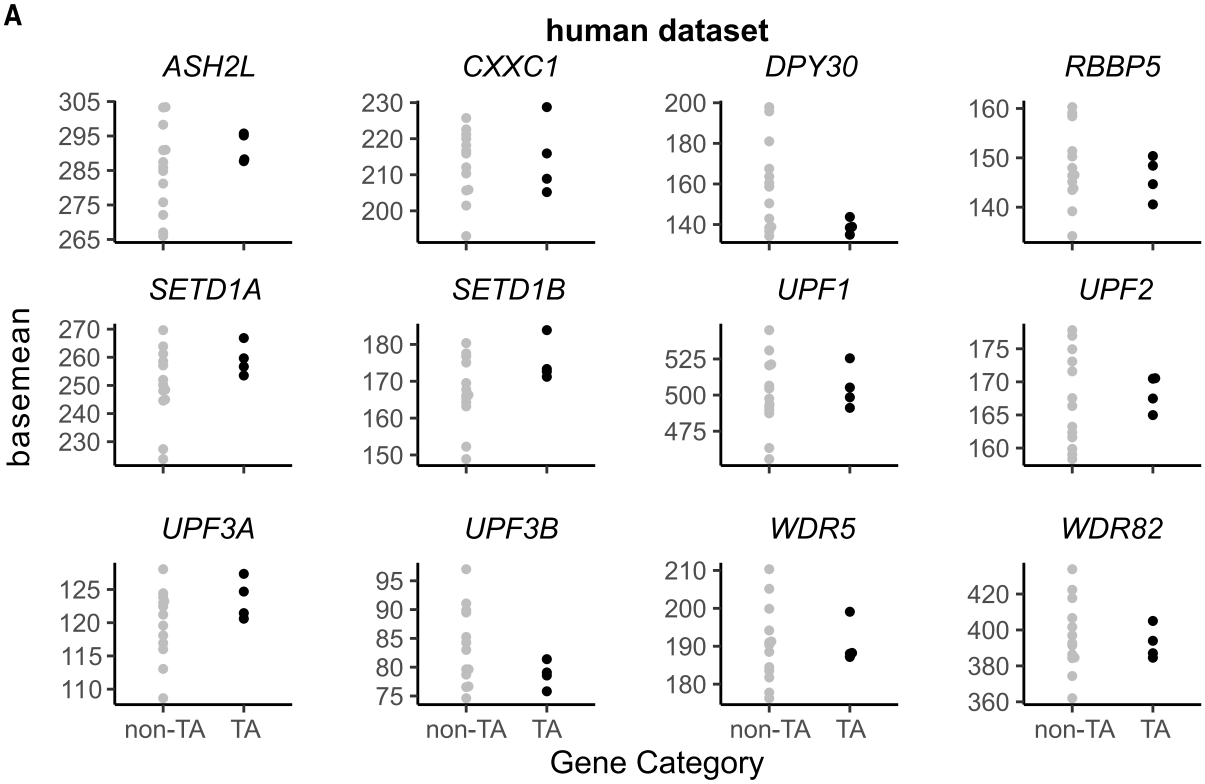


B

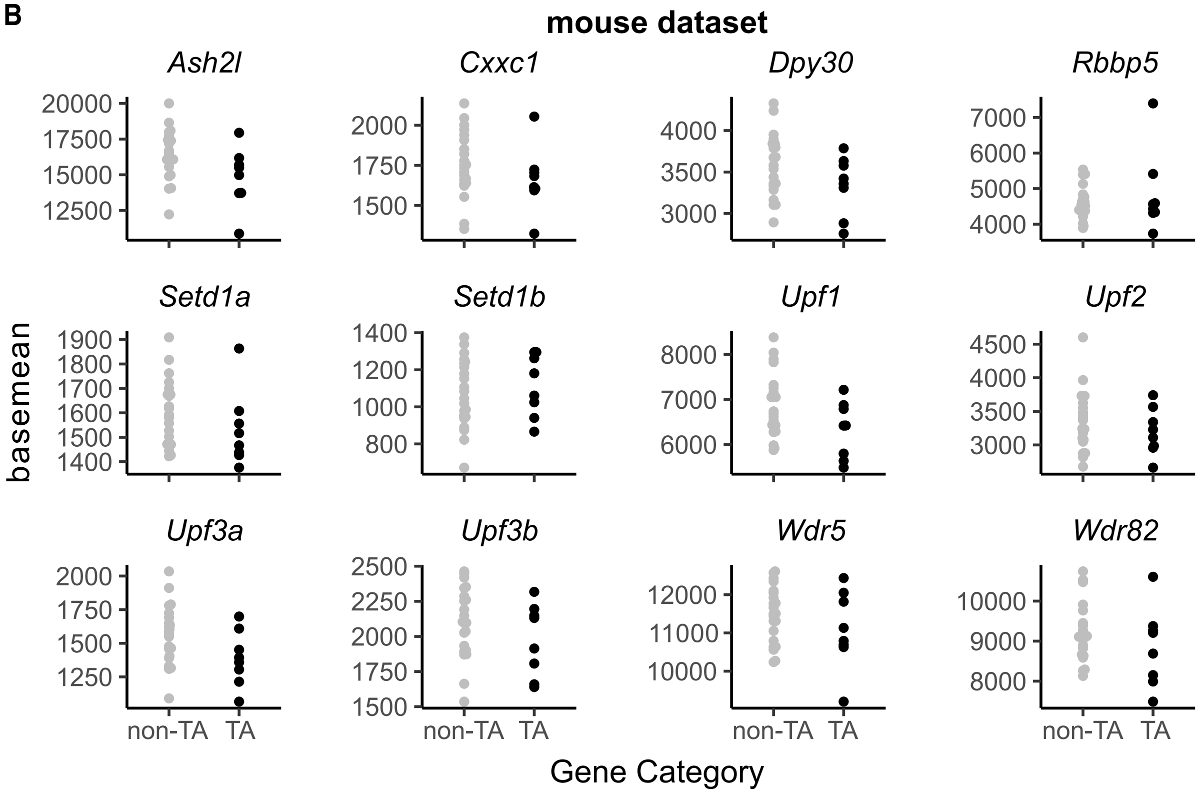


**Supplementary Figure 2**

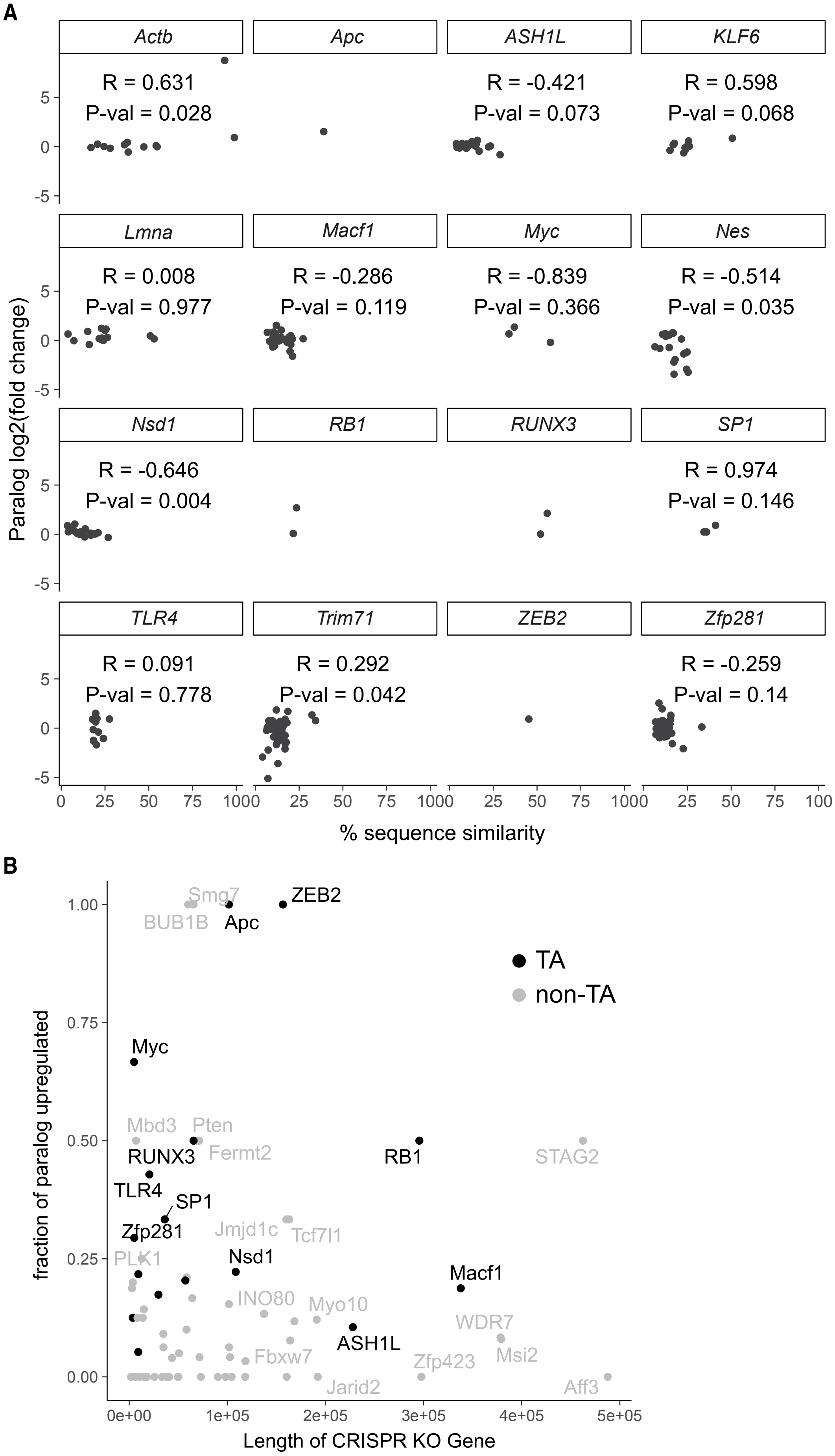


**Supplementary Figure 3**

**A**

| Parameter | Description |
|---|---|
| $r_{on}$ | Activation rate of an allele |
| $r_{off}$ | Inactivation rate of an allele |
| $r_{prod}$ | Production rate of mRNA from an active allele |
| $r_{deg}$ | Degradation rate of mRNA |
| $r_{add}^{NITC}$ | Additional activation of an allele (A or A') upon nonsense-induced transcriptional compensation caused by product $A_{nonsense}$ |
| $r_{add}^{A,B}$, $r_{add}^{A\prime,B}$ | Activation of B by A or A' to one of two respective B active states specified in the model |
| $d$ | Factor by which the mRNA production rate of B is lower in the A'-directed B active state than in A-directed B active state |
| $n$ | Hill coefficient |
| $k$ | Dissociation constant of the Hill function |

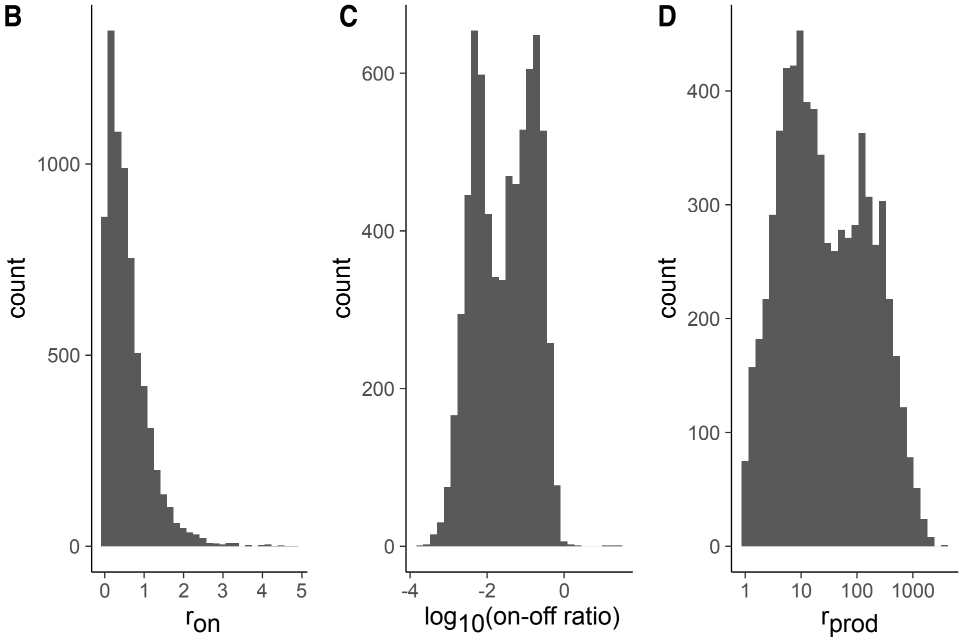


**E**

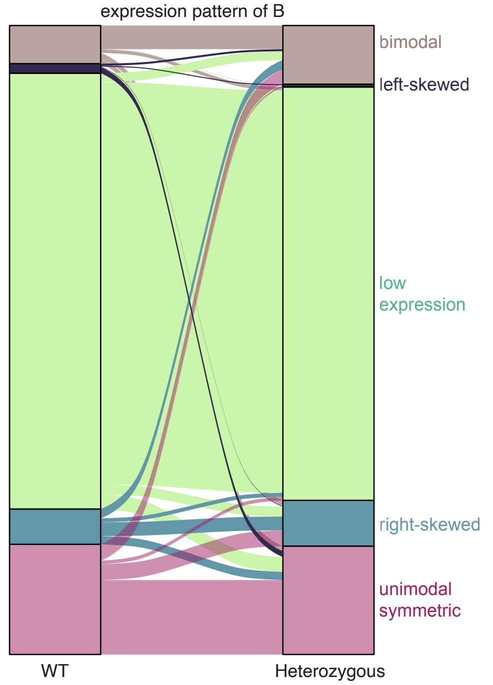


**Supplementary Figure 4**

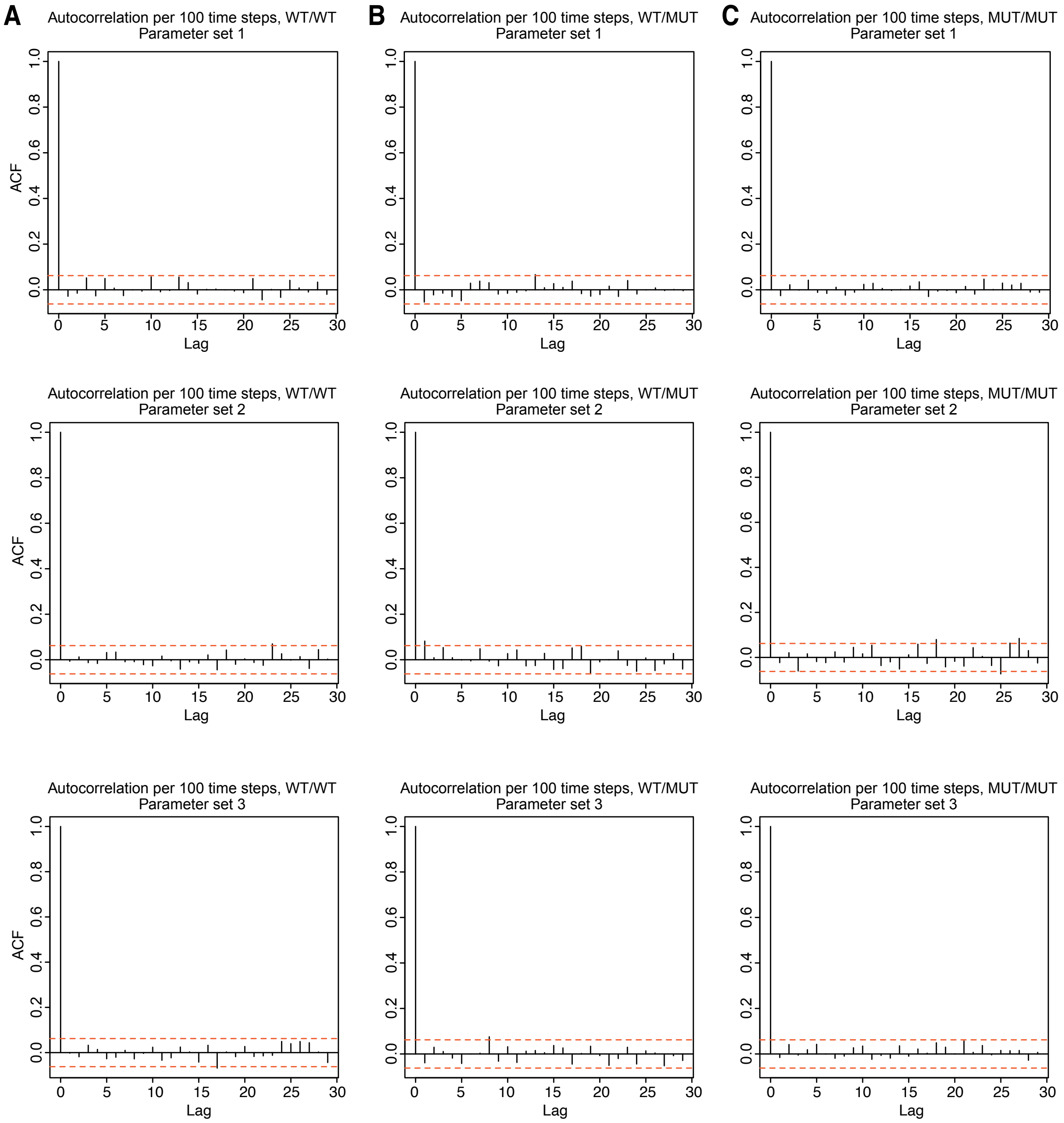


**Supplementary Figure 5**

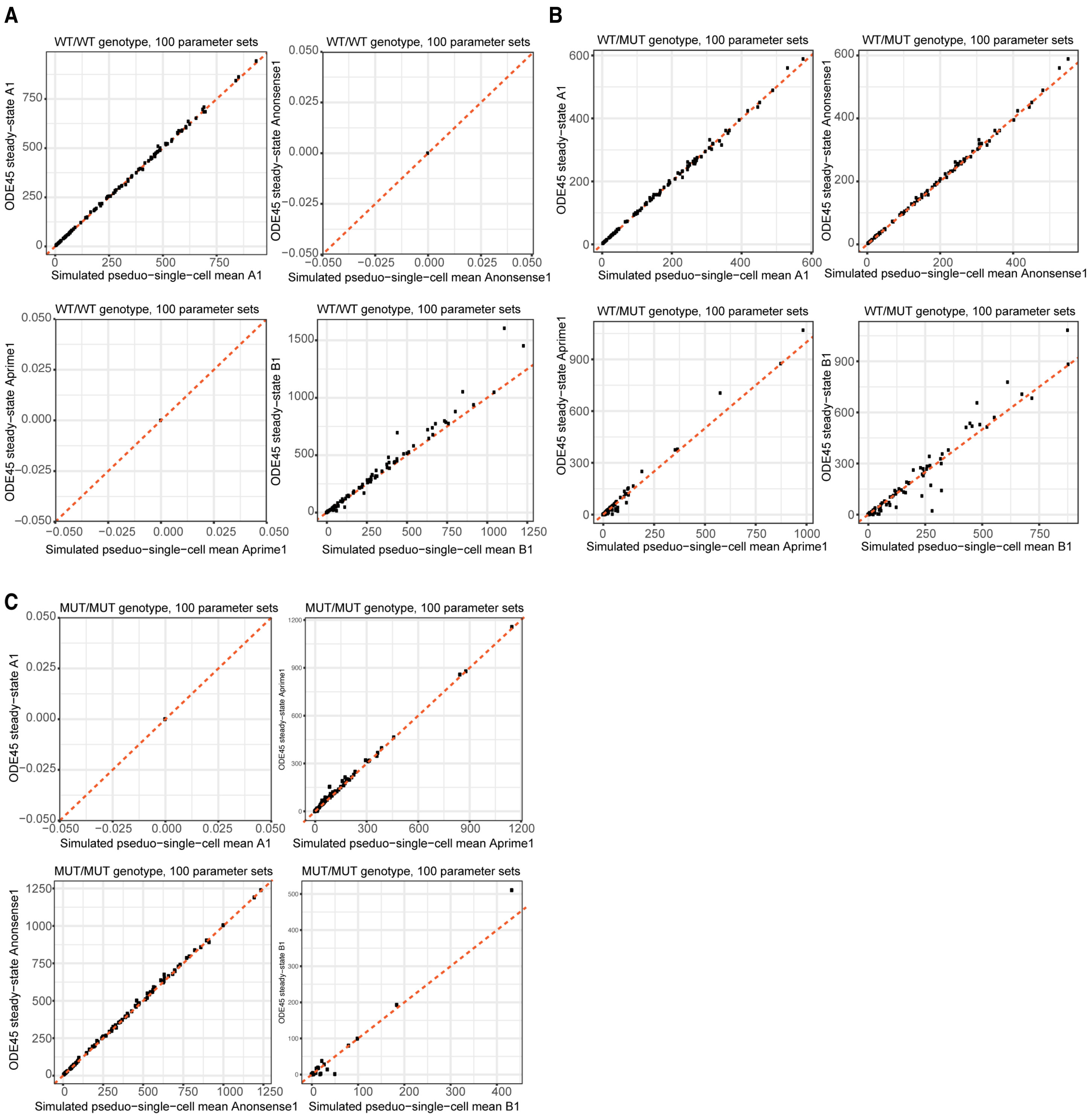


**Supplementary Figure 6**

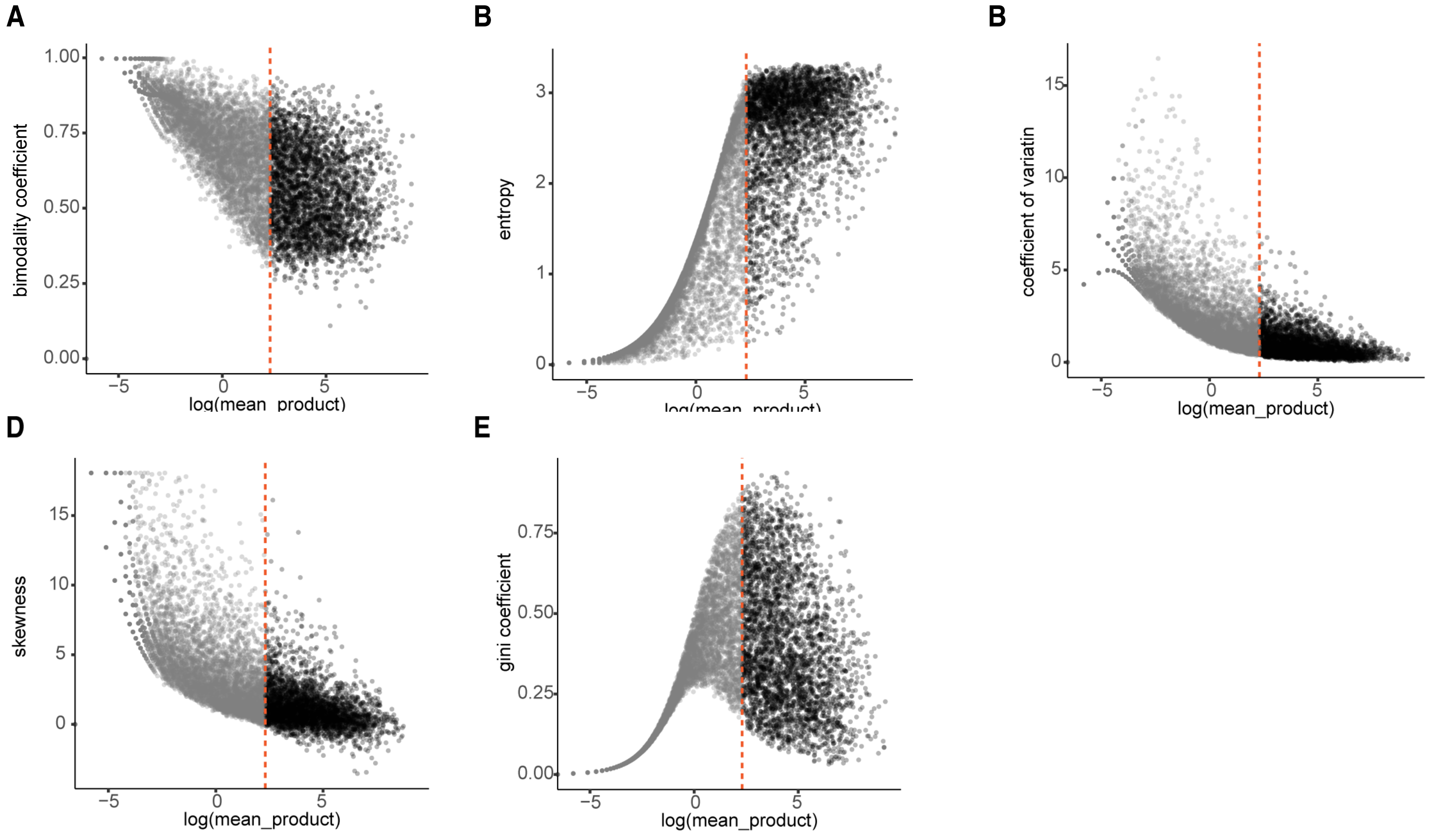


**Supplementary Figure 7**

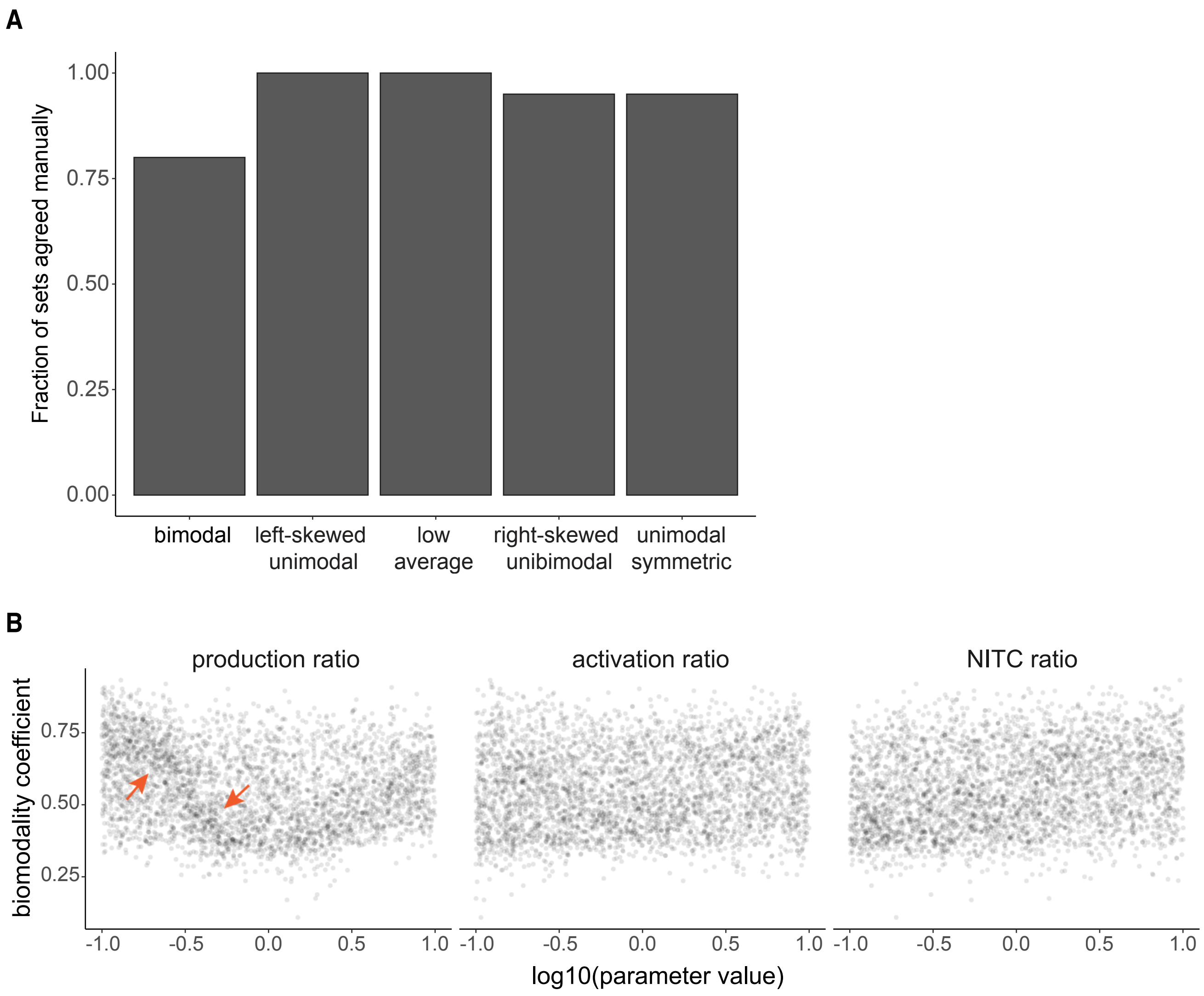


**Supplementary Figure 8**

**A**

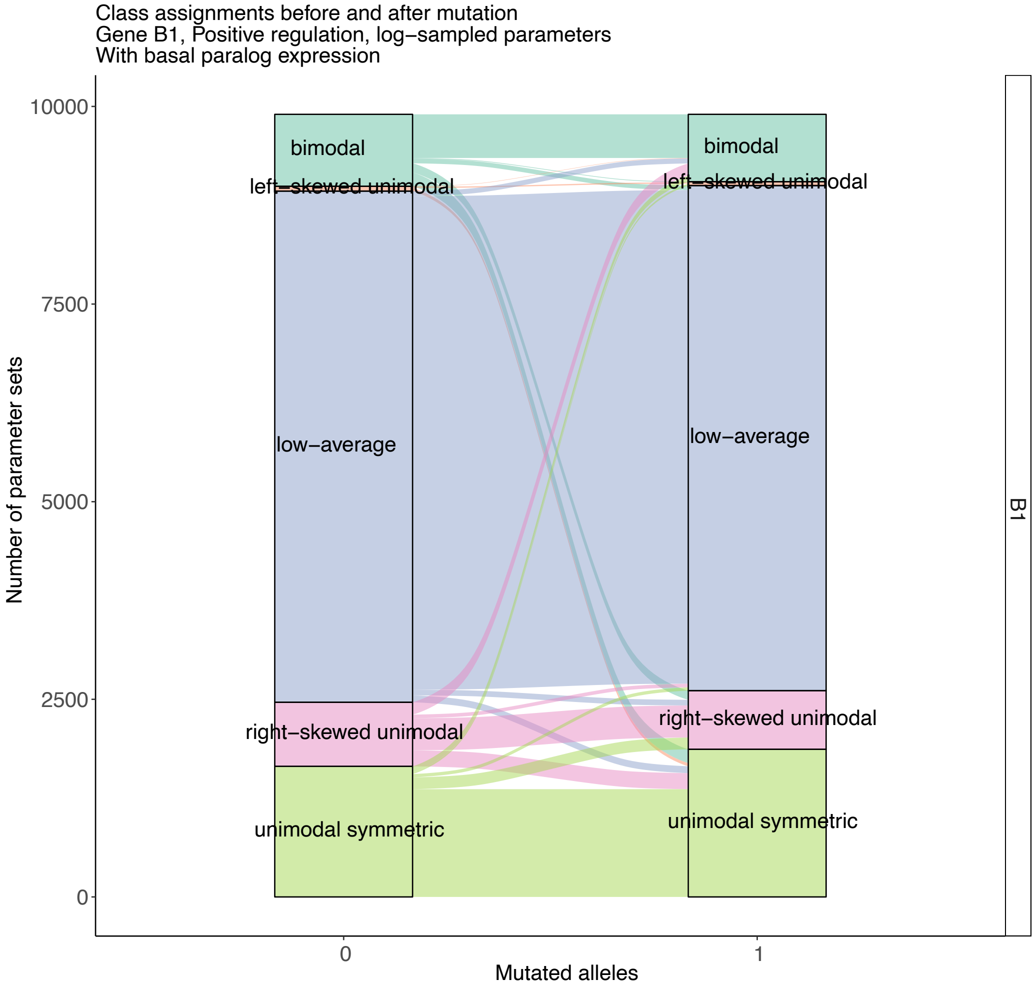


**Supplementary Figure 9**

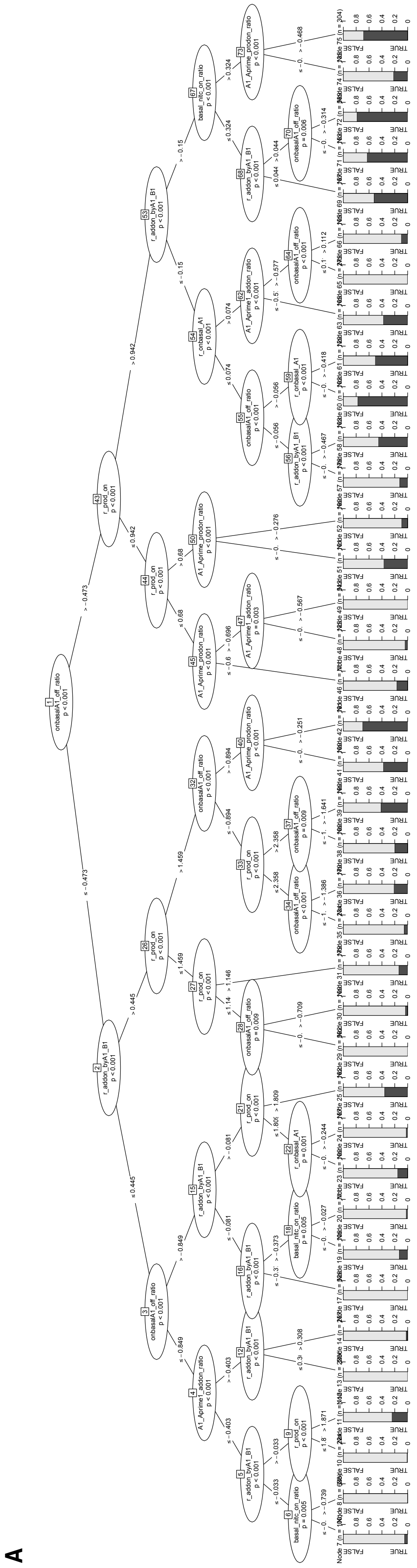


**Supplementary Figure 10**

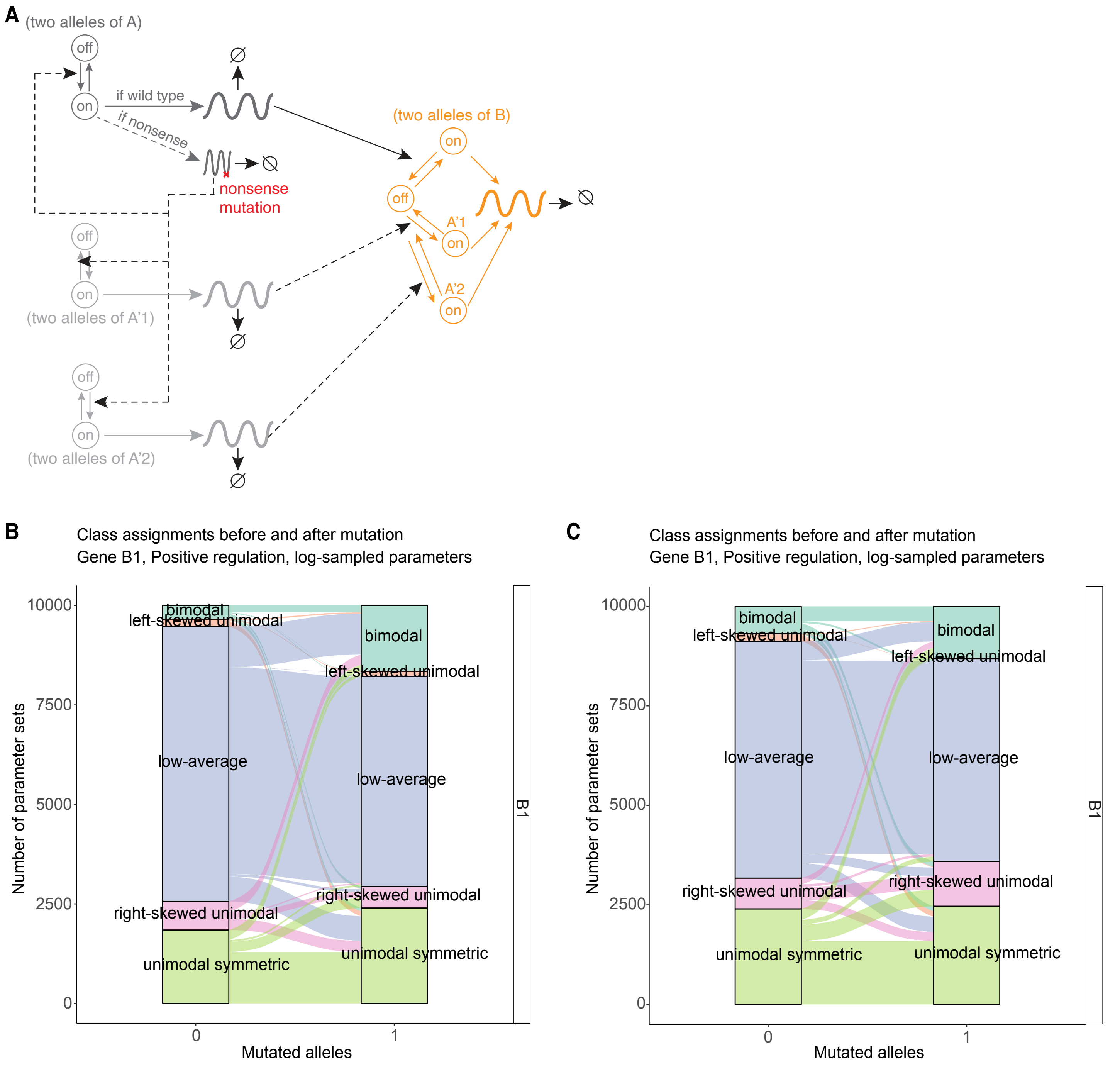


**Supplementary Figure 11**

A

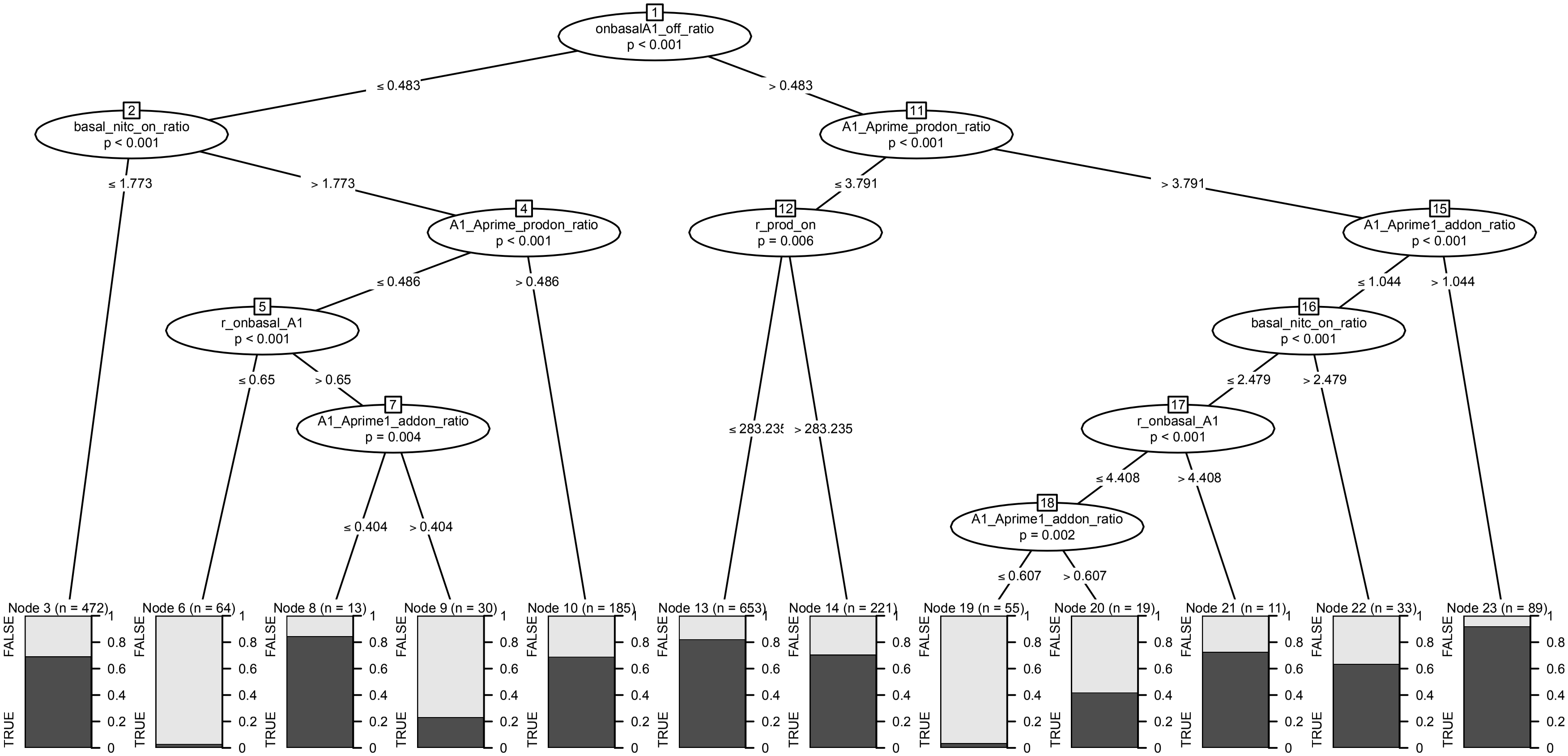


Supplementary Figure 12

A

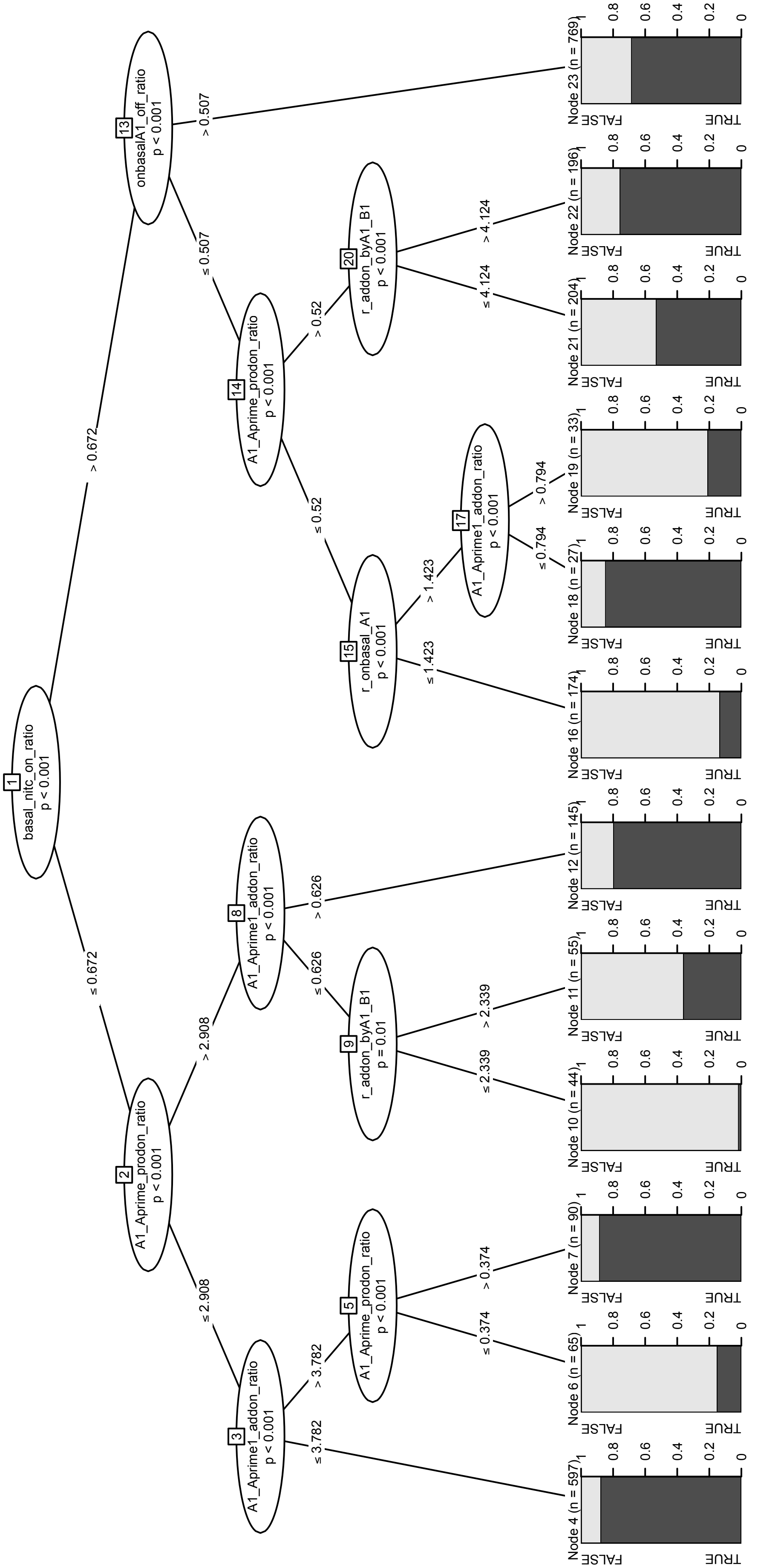


**Supplementary Figure 13**

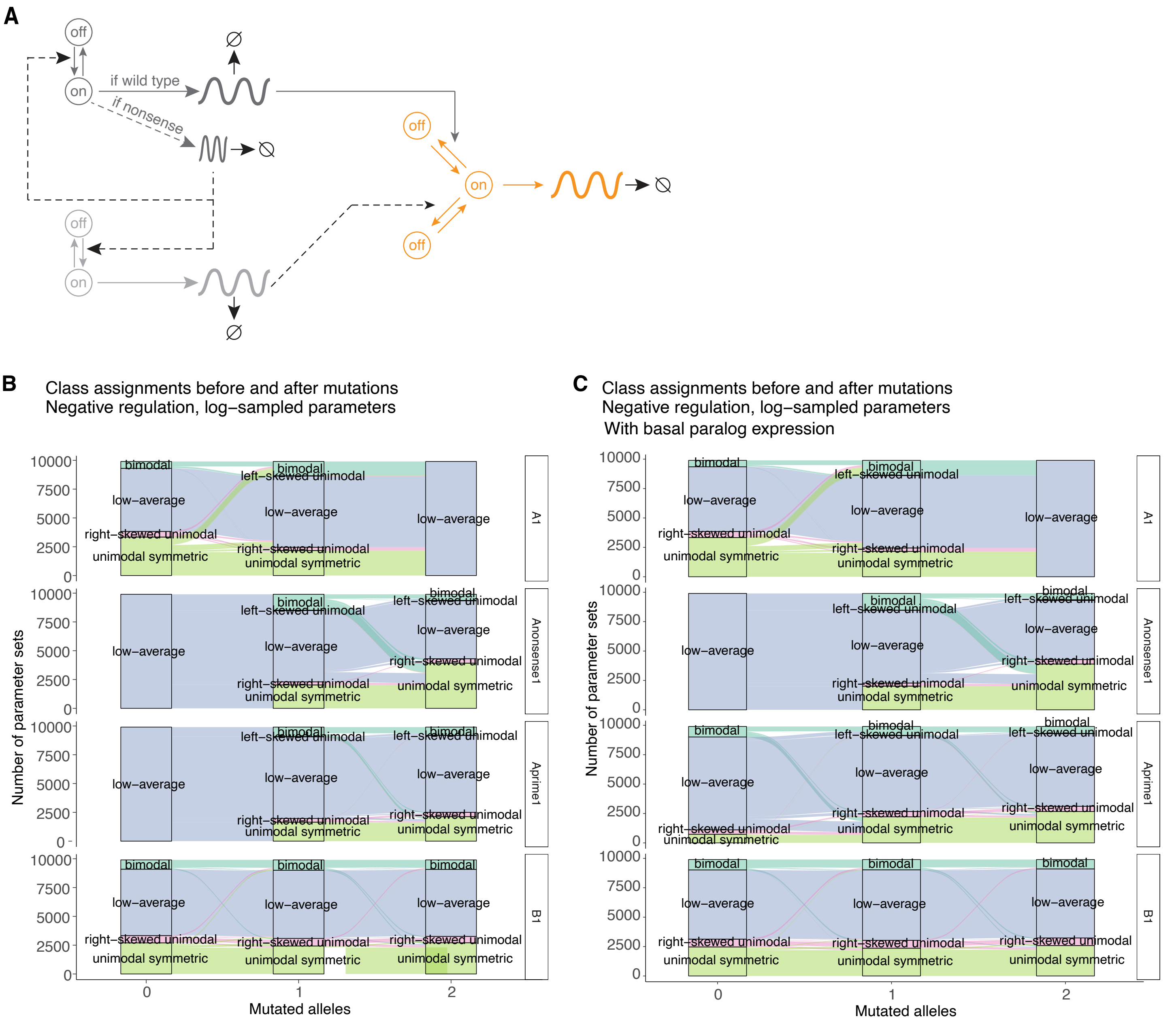


**Supplementary Figure 14**

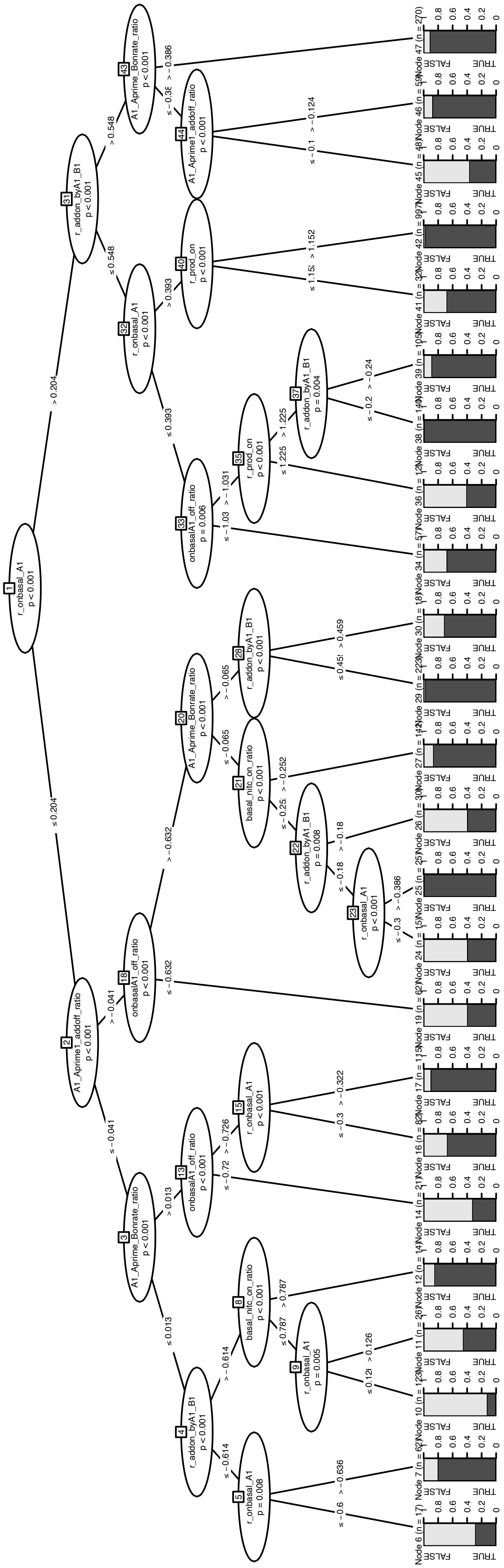

A

**Supplementary Figure 15**

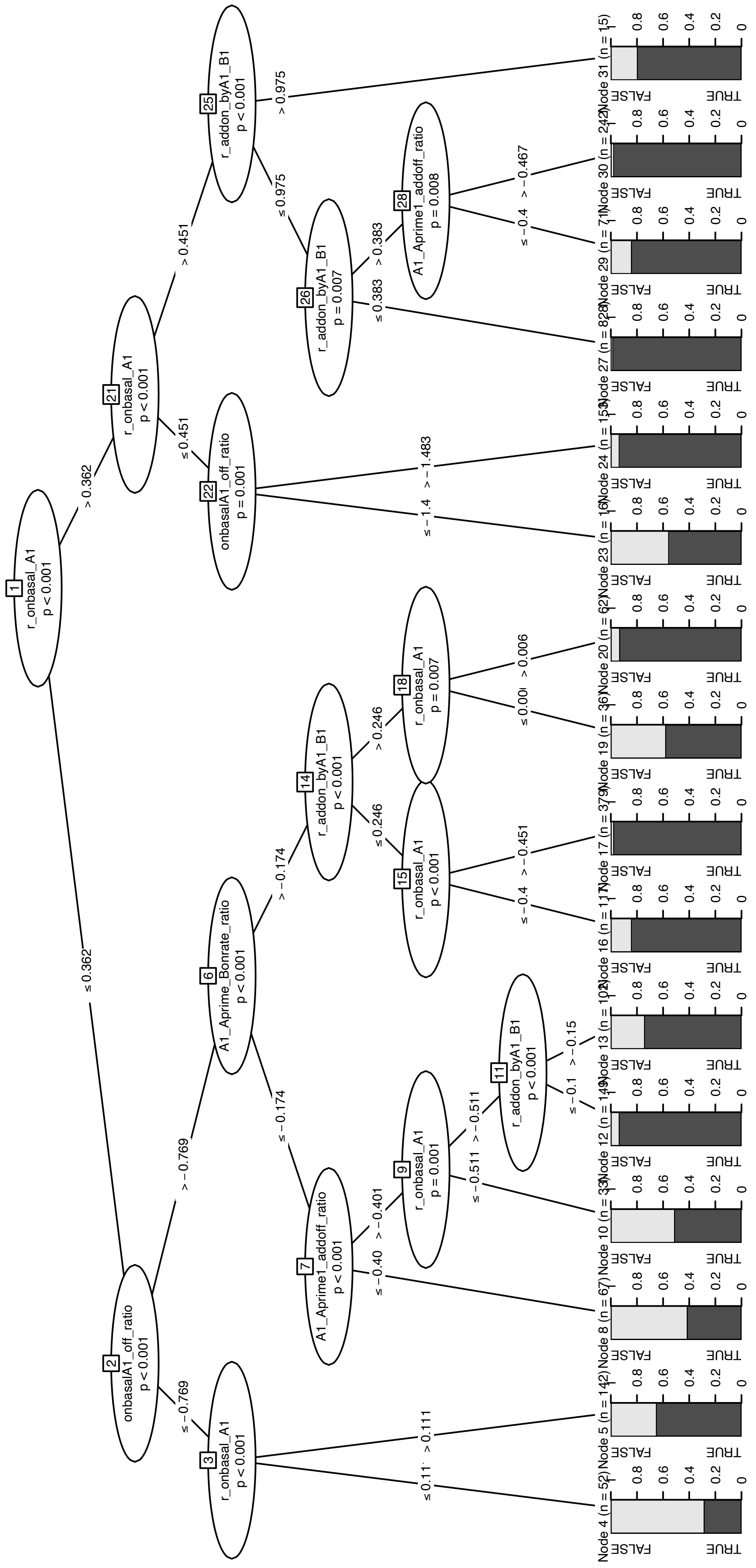


A

**Supplementary Figure 16**
